# MPZ-T124M mouse model replicates human axonopathy and suggest alteration in axo-glia communication

**DOI:** 10.1101/2022.05.09.491190

**Authors:** Ghjuvan’Ghjacumu Shackleford, Leandro N. Marziali, Yo Sasaki, Nadav I. Weinstock, Alexander M. Rossor, Nicholas J. Silvestri, Emma R. Wilson, Edward Hurley, Grahame J. Kidd, Senthilvelan Manohar, Dalian Ding, Richard J. Salvi, M. Laura Feltri, Maurizio D’Antonio, Lawrence Wrabetz

## Abstract

Myelin is essential for rapid nerve impulse propagation and axon protection. Accordingly, defects in myelination or myelin maintenance lead to secondary axonal damage and subsequent degeneration. Studies utilizing genetic (CNPase-, MAG-, and PLP-null mice) and naturally occurring neuropathy models suggest that myelinating glia also support axons independently from myelin. Myelin protein zero (MPZ or P0), which is expressed only by Schwann cells, is critical for myelin formation and maintenance in the peripheral nervous system. Many mutations in *MPZ* are associated with demyelinating neuropathies (Charcot-Marie-Tooth disease type 1B [CMT1B]). Surprisingly, the substitution of threonine by methionine at position 124 of P0 (P0T124M) causes axonal neuropathy (CMT2J) with little to no myelin damage. This disease provides an excellent paradigm to understand how myelinating glia support axons independently from myelin. To study this, we generated targeted knock-in P0T124M mutant mice, a genetically authentic model of T124M-CMT2J neuropathy. Similar to patients, these mice develop axonopathy between 2 and 12 months of age, characterized by impaired motor performance, normal nerve conduction velocities but reduced compound motor action potential amplitudes, and axonal damage with only minor compact myelin modifications. Mechanistically, we detected metabolic changes that could lead to axonal degeneration, and prominent alterations in non-compact myelin domains such as paranodes, Schmidt-Lanterman incisures, and gap junctions, implicated in Schwann cell-axon communication and axonal metabolic support. Finally, we document perturbed mitochondrial size and distribution along P0T124M axons suggesting altered axonal transport. Our data suggest that Schwann cells in P0T124M mutant mice cannot provide axons with sufficient trophic support, leading to reduced ATP biosynthesis and axonopathy. In conclusion, the P0T124M mouse model faithfully reproduces the human neuropathy and represents a unique tool for identifying the molecular basis for glial support of axons.

## Introduction

In the peripheral nervous system (PNS), Schwann cells (SC) make myelin by wrapping their plasma membranes around axons. Myelin is a multilamellar lipid-rich structure containing a set of proteins (in the PNS: mostly myelin protein zero [MPZ, P0], peripheral myelin protein 22 [PMP22], and myelin basic protein [MBP]) essential for its compaction. Among these structural proteins, P0, encoded by the *MPZ* gene, is the most abundant, accounting for 45% of the total protein expressed in PNS myelin^1^. P0 is a single-pass transmembrane glycoprotein with an immunoglobulin-like fold in its extracellular domain that is expressed exclusively by SC. Crystallography of P0 and X-ray diffraction analysis of myelin suggest that P0 forms homotetramers interacting *in trans* to hold together adjacent wraps of myelin membrane^2^. As evidence of the crucial role of P0 in myelination and myelin compaction^3^, *Mpz* deficient mice produce few myelin layers that are poorly compacted^4^.

Charcot-Marie-Tooth (CMT) neuropathies are the most common inherited neurological disorders, affecting 1/2500 people^5^. CMT neuropathies are caused by alterations in over 80 genes encompassing approximately 1,000 independent mutations^6^. Based on nerve conduction velocity (NCV), CMT are mainly classified as demyelinating CMT type 1 (CMT1) (NCV<38 m/s) or axonal CMT type 2 (CMT2) (NCV>38 m/s) hereditary neuropathies. CMT1B, represents a special challenge because it is caused by more than 200 diverse mutations in *MPZ,* resulting in different toxic gain of function mechanisms and various related clinical phenotypes^7, 8^. Initially, the majority of P0 mutations identified were associated with demyelinating neuropathies. CMT1B patients typically present as early onset disease with extremely slow NCV (<10-20 m/s). Surprisingly, since P0 is only expressed in SC but not in neurons, several mutations in P0^9–12^, such as the substitution of threonine by methionine at position 124 (P0T124M), were shown to cause an axonal neuropathy referenced as CMT2J. After around 40 years of age, patients with T124M-CMT2J begin experiencing symptoms, such as a lancinating pain and fast-progressive weakness of the lower limbs. NCVs vary widely among T124M-CMT2J patients and can be normal, slightly reduced similar to that seen in CMT2, or as slow as in CMT1 but never as low as in CMT1B. However, the amplitudes of motor evoked potentials are strikingly reduced in patients carrying the P0T124M mutation. Sural nerve biopsy samples exhibit regenerative clusters and a marked reduction of myelinated axons, but with little to no myelin damage. Moreover, T124M-CMT2J patients exhibit early signs of sensory abnormalities, such as pupillary abnormalities and hearing loss^13–15^. Beside its myelin producing function, these observations suggest either that P0 in SC contributes to axonal support or that SC expressing P0T124M produce deleterious signals contributing to axonal degeneration.

Axonal degeneration is a common endpoint of peripheral neuropathies^16^ and understanding the glial processes that contribute to axon protection and degeneration is considered the key to cure neuropathies. Yet, this topic is poorly understood. Axonal degeneration is uncoupled from demyelination in CMT2J, providing a unique opportunity to understand how myelinating glia support axons independently of myelin. We generated an authentic mouse model of CMT2J carrying the P0T124M mutation, representing, to the best of our knowledge, the first animal model for CMT2J disease and the first model of human axonal neuropathy caused by a mutation in a protein of compact myelin. Our findings show that the P0T124M mouse model closely recapitulates the axonopathy and clinical aspects observed in CMT2J patients. Alterations in non-compact myelin, metabolic changes, and mitochondria dysfunction in P0T124M mutants suggest that axonal loss is the result of a defect in SC-to-axon communication. Our results highlight potential mechanisms of how a mutation in P0 causes axonal degeneration without demyelination and underlie the fundamental role of SC in axonal support.

## Materials and methods

### P0T124M transgenic mice

All experiments performed on mice were conducted in accordance with experimental protocols approved by the Institutional Animal Care and Use Committees of Roswell Park, University at Buffalo, and San Raffaele Scientific Institute. The P0T124M mutation was targeted to a single *Mpz* allele by homologous recombination as described by Saporta *et al.*^17^ for the generation of P0R98C mice. The mutation was introduced into *Mpz* exon 3 of a 129S2 genomic clone by site-directed mutagenesis and confirmed by sequence analysis. Fragments of this clone were ligated into a construct containing the neomycin resistance gene (*neoR*) flanked by loxP sites (**Fig. S1A**). The T124MneoLP construct was electroporated into TBV2 (129S2 strain) embryonic stem cells as described previously by Nodari *et al*^18^. After confirming homologous recombination by Southern blot analysis with probes recognizing the sequence flanking either the long or the short arm of the construct, positive embryonic stem cell clones were injected into blastocysts of host wild-type animals to obtain chimeras. One chimera transgene was germline transmitted, and the mouse harboring the transgene was crossed with CMV-Cre mice (JAX#006054) to excise the *neoR* cassette (**supplementary Fig. 1A**). To confirm that the mutation was present as expected, RNA from the sciatic nerves of P0T124M mice was isolated and reverse transcribed. The full coding sequence of *Mpz* cDNA was amplified by PCR (forward (F), 5′-ATGGCTCCCGGGGCTCCC-3′; reverse (R), 5′- CTATTTTCTTATCCTTGCGAG-3′), cloned, and sequenced. Genotyping was performed by PCR analysis. The P0T124M mutation introduced an extra NSP1 restriction enzyme digestion site in *Mpz* exon 3 (**supplementary Fig. 1B and C**). PCR primers MpzEx3F (5′- CGATGAGGTGGGGGCCTTCAA-3′) and MpzEx3R (5′-ATAGAGCGTGACCTGAGAGG-3′) generate a 169 base pairs (bp) amplimer. After NSP1 digestion, the wild-type allele migrates at 114 bp + 55 bp, whereas the T124M allele migrates at 106 bp + 8 bp + 55 bp. The 8-bp band was not detected on the PCR gel (**supplementary Fig. 1D and E)**.

**Figure 1:**
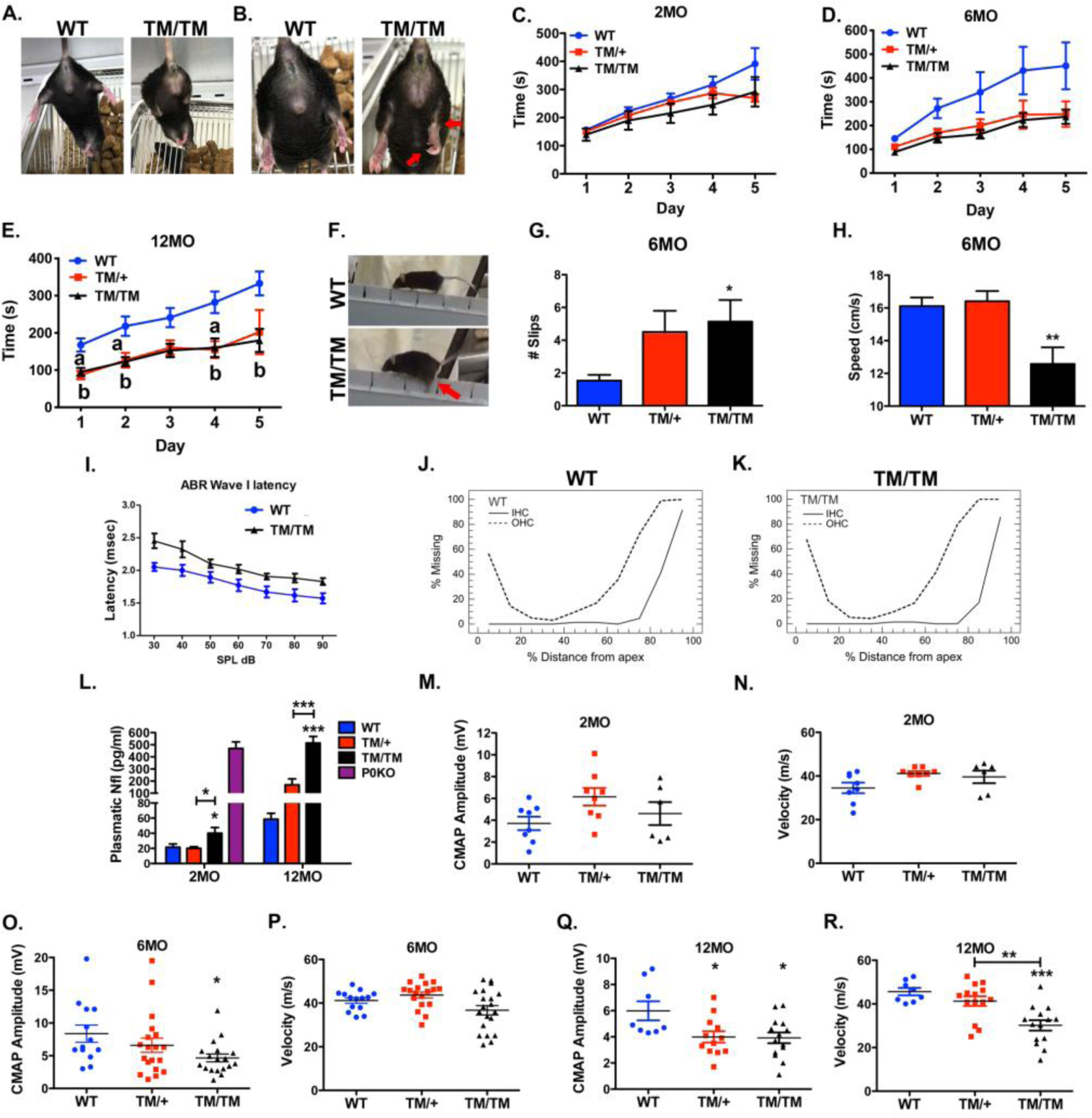
Clinical impairments in TM mice. (**A and B**) External signs of neuropathy in 12- month-old P0T124M homozygous (TM/TM) mice. (**A**) Representative picture of clasping behavior. (**B**) Clawed hind paws and Achilles’ tendon retraction are indicated by arrows. (**C to G**) Locomotion is impaired in TM mice. Accelerating rotarod test at 2 (**C**), 6 (**D**), and 12 (**E**) months of age. No differences were observed at 2 months of age. Compared to that of wild-type (WT) mice, the performance of TM mice was worse at 6 months of age and significantly altered at 12. *n* (animals) ≥ 4 per genotype. Two-way ANOVA with Dunnett’s *post hoc* test analysis comparing WT with TM/+ mice (*^a^p < 0.05*) and WT with TM/TM mice (*^b^p < 0.05*). (**F to H**) Beam walking test at 6 months of age. (**F**) Representative pictures of beam walking test for WT and TM/TM mice. Arrow indicates a slip. Quantification of slips (**G**) and speed (**H**). *n* (animals) ≥ 5 per genotype. One-way ANOVA with Dunnett’s *post hoc* test analysis comparing WT with TM/+ mice and WT with TM/TM mice. \**p < 0.05*, \*\**p < 0.01*. (**I to K**) Auditory neuropathy in TM mice. (**I**) ABR wave I latency measurements in WT and TM/TM mice at 11 months of age. Cochleograms of WT (**J**) and TM/TM (**K**) mice at 12 months of age. *n* (animals) ≥ 4 per genotype. (**L**) Plasmatic neurofilament light (Nfl) concentrations in WT, TM/+, and TM/TM mice at 2 and 12 months of age. Plasma from 4-month-old P0 null mice was used as a positive control. *n* (animals) ≥ 5 per genotype. (**M to R**) Electrophysiological analysis. Amplitudes of compound muscle action potentials (CMAPs) and nerve conduction velocities (NCVs) were measured at 2 (**M and N**), 6 (**O and P**), and 12 (**Q and R**) months of age. *n* (nerves) ≥ 6 per genotype. \**p < 0.05*, \*\**p < 0.01*, \*\*\**p < 0.001* by Tukey’s *post hoc* tests (**L to S**) after one-way ANOVA. Graphs indicate means ± SEMs.

### Statistical analyses

Experiments were not randomized, but data collection and analysis were performed blindly to the conditions of the experiments. No statistical methods were used to predetermine sample sizes, but the sample sizes are similar to those generally employed in the field. All statistical analyses were conducted using GraphPad Prism 6.01. To determine the significance between genotypes, Student’s *t* tests and one-way and two-way ANOVAs with Tukey’s and Dunnett’s comparison tests were used. A *p* value of ≤0.05 was considered statistically significant.

### Data availability

All data in this report are available to all readers. The reagents used in this study, as well as further information, are available upon request from M.L.F.

Detailed materials and methods are reported in the supplementary material.

## Results

### T124M mice display progressive motor defects

Using homologous recombination, we engineered a genetically authentic T124M-CMT2J mouse model (**supplementary Fig. 1A, B, and C**). Like CMT2J patients, the mice harboring the P0T124M mutation (referred to as TM mice) exhibit signs of late-onset, progressive peripheral neuropathy. Starting at 6 months of age, and more obviously at 12 months, we noticed limb clasping behavior, clawed hind paws, and Achilles’ tendon retraction (**Fig. 1A and B**). These neurological abnormalities were not fully penetrant, reflecting the heterogeneity of symptoms observed in T124M-CMT2J patients, and were more prominent in TM homozygous (TM/TM) mice than in heterozygous (TM/+) mice.

We tested the motor performance of TM mice in the rotarod and beam walking tests and found no differences from wild-type (WT) mice at 2 months of age (**Fig. 1C**). However, at 6 months of age (**Fig. 1D**), we noticed a trend towards a reduced motor capacity in TM mice; by 12 months (**Fig. 1E**), TM mice remained on the accelerating rotarod half as long as WT mice. Results from the beam walking test further demonstrated motor impairments (**Fig. 1F and supplementary movies 1, 2 and 3**). At 6 months of age, TM/+ and TM/TM mice exhibited more foot slips (**Fig. 1F and G**), and TM/TM mice were 25% slower than the WT mice in crossing the rod (**Fig. 1H**). Thus, TM mice develop a peripheral neuropathy and display locomotion impairment.

### Hearing loss in T124M mice

Hearing loss is one of the most consistently observed symptoms in T124M-CMT2J patients^13, 14^. To determine if hearing loss also occurs in our mouse model, we used click stimuli to obtain auditory brainstem evoked potential recordings (ABR). The ABR trace consists of five major peaks or waves occurring in the first 5-7 ms following stimulus onset. Wave I, which has a latency of approximately 1.8 ms at high intensities, reflects neural activity of the auditory nerve and subsequent peaks represent to a first approximation synchronized neural responses from successfully more proximal regions in the auditory brainstem (i.e. cochlear nuclei, superior olive, lateral lemniscus, and inferior colliculus). Because only the distal part of the auditory nerve is myelinated by SC, while more central structures in the CNS are myelinated by oligodendrocytes, we focused our attention on ABR wave I. By 11 months of age, the click-evoked latency of ABR wave I was markedly increased at all stimulus intensities in TM/TM mice compared to WT mice (**Fig. 1I**). The increase in latency is suggestive of a neural conduction delay in the auditory nerve similar to that observed in T124M-CMT2J patients. To determine if this latency prolongation was caused by loss of sensory hair cells, we performed a cochleogram assay to assess the percentage loss of outer hair cells (OHC) and inner hair cells (IHC) from the base to the apex of the cochlea. Mean cochleogram revealed losses of OHC and to a lesser extent IHC in the basal half of the cochlea of both TM/TM and WT mice. Because there were no significant differences in the magnitude of OHC and IHC lesions between WT and TM/TM mice, the longer wave I latencies in TM/TM mice compared to WT mice are presumably caused by defect on peripheral auditory nerve fibers (**Fig. 1J and K**). Abnormal ABRs are characteristic of the auditory neuropathy observed in late-onset CMT1B patients harboring the P0Y145S mutation^19^, patients with *PMP22* mutations^20, 21^, and those with *GJB1* (gap junction beta 1) mutations^20, 22^, suggesting that hearing impairment in those with the P0T124M mutation may be due in part to degeneration of the distal part of the auditory nerve.

### Level of plasmatic Neurofilament light is increased in T124M mice

Plasmatic neurofilament light (pNfl) is emerging as a biomarker for a variety of neurological diseases and positively correlates with the severity of peripheral neuropathies^23^. Remarkably, pNfl concentrations were increased 2-fold in TM/TM mice at 2 months of age and 8-fold at 12 months of age. In TM/+ mice, pNfl concentrations were trending toward an increase (3-fold) at 12 months of age (**Fig. 1L**). These data suggest that pNfl level could also serve as a biomarker in T124M-CMT2J patients and may correlate with disease progression.

### Electrophysiological alterations in T124M mice

Electrophysiological analyses showed no differences between TM and WT mice at 2 months of age (**Fig. 1M and 1N**). At 6 months of age, there was a trend toward reduced compound muscle action potential (CMAP) amplitude in TM/+ mice and a significantly reduced CMAP amplitude in TM/TM mice (**Fig. 1O**) but normal NCVs (**Fig. 1P**). At 12 months, both TM/TM and TM/+ mice exhibited significantly reduced CMAP amplitudes (**Fig. 1Q**), consistent with the observed neuromuscular impairment, but only TM/TM mice had significantly lower NCVs (**Fig. 1R**). Altogether, these results show that the P0T124M mutation causes a peripheral neuropathy with negative functional impact on axons before myelin.

### T124M mice develop a progressive axonopathy with subtle myelin defects

We next studied the consequence of P0T124M mutation on nerve morphology (**Fig. 2 and supplementary Fig. 2**). Because P0 is the most aboundant myelin protein and because P0 mutations are responsible for hypomyelinating, dysmyelinating, and demyelinating CMT1B neuropathies, we first studied the ultrastructure of myelin sheaths in sciatic nerves from TM mice (**Fig. 2A to D**). At 2 and 6 months of age, there were no significant differences in myelin thickness between WT and TM mice. By contrast, we detected thicker myelin sheaths, reflected by significantly lower g-ratios, in 12- and 18-month-old TM/TM and TM/+ mice (**Fig. 2E**). The increase in myelin thickness was predominantly observed for small-diameter axons (**Fig. 2F**). To determine whether the increased thickness is attributable to a defect in myelin compaction, we measured myelin periodicity. There was a slight but not significant, increase of myelin periodicity in TM mice (**Fig. 2G and H**). Consistent with this result, the expression of major myelin proteins (myelin-associated glycoprotein [MAG], PMP22, 2’,3’-cyclic nucleotide 3’-phosphodiesterase and [CNPase]) was not affected in TM sciatic nerve at 2 and 6 months of age. Similar results were obtained at 18 months of age, except for a decrease in CNPase expression (**supplementary Fig. 3**). These results corroborate the observations of samples from T124M-CMT2J patients, in which compact myelin was not or only mildly altered and was well organized^14, 15^.

**Figure 2:**
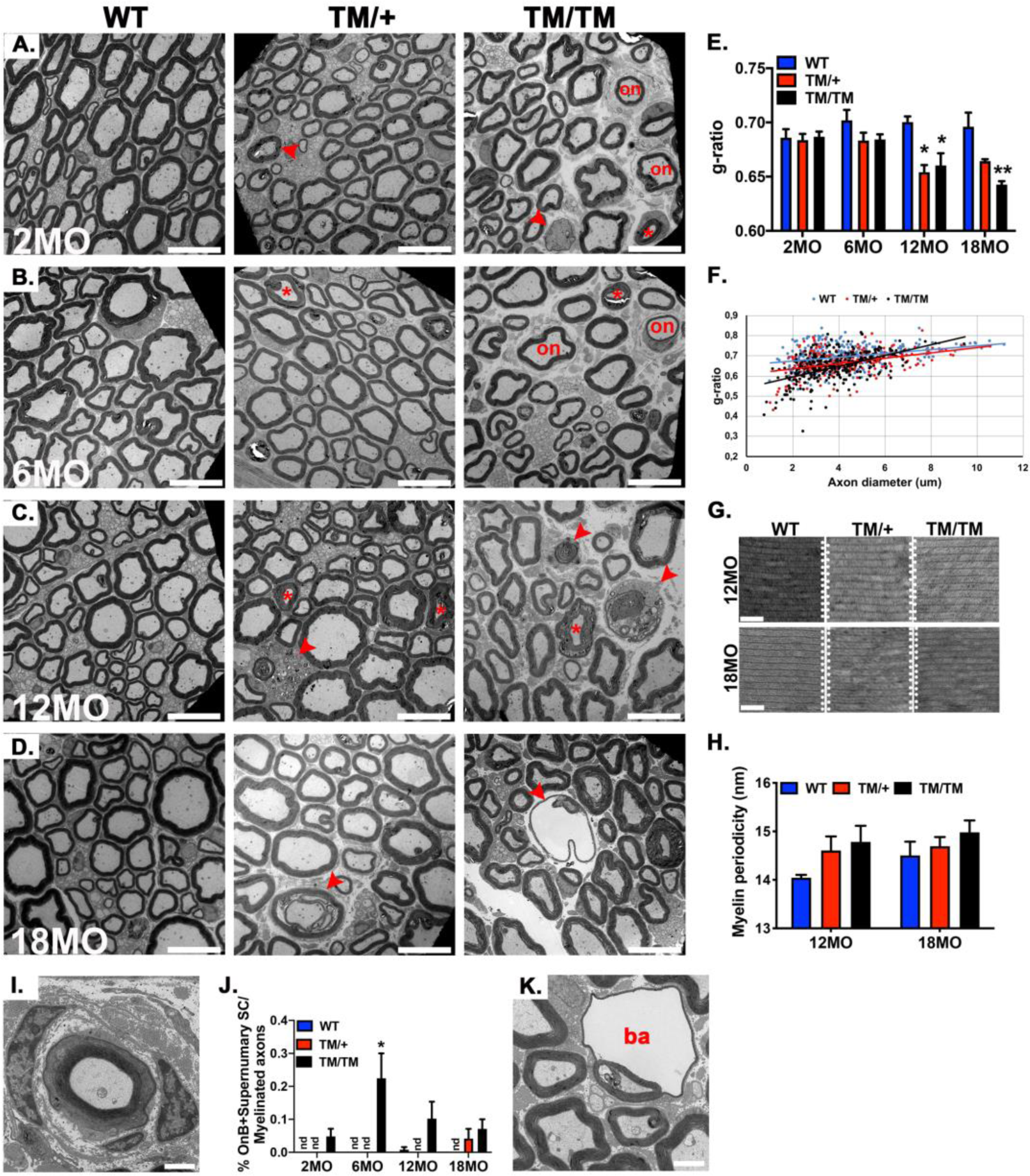
Morphological analysis of TM nerves. (**A to D**) Representative electron micrographs of wild-type (WT), P0T124M heterozygous (TM/+), and homozygous (TM/TM) sciatic nerve cross sections at 2 (**A**), 6 (**B**), 12 (**C**), and 18 (**D**) months of age. Arrowheads indicate degenerative figures, asterisks indicate SLI, and “on” indicate onion bulbs and supernumerary SC. Scale bars: 10 μm. (**E**) g-ratio analysis shows hypermyelination of TM fibers starting at 12 months of age. (**F**) g-ratio as a function of axonal diameter shows preferential hypermyelination of small fibers in TM mice compared to that in wild-type (WT) mice. (**G**) Ultrastructural analysis of periodicity shows that myelin sheaths are well compacted in TM mice. Scale bars: 50 μm. (**H**) Myelin periodicity was slightly, but not significantly, increased in TM mice. (**I**) Representative electron micrograph of an onion bulb. (**J**) Onion bulbs and supernumerary SC were rare in TM/TM mice. They represented 0.2% of myelinated fibers in 6-month-old mice and did not progress over time. (**K**) Representative electron micrograph of a myelin balloon formed by fluid inside the intraperiodic line of the myelin sheath. Note how the axon is compressed. **I** and **N** scale bars: 2 μm. *n* (animals) ≥ 3 per genotype. \**p < 0.05*, \*\**p < 0.01*, by multiple-comparisons Tukey’s *post hoc* tests after one-way ANOVA. Graphs indicate means ± SEMs.

**Figure 3:**
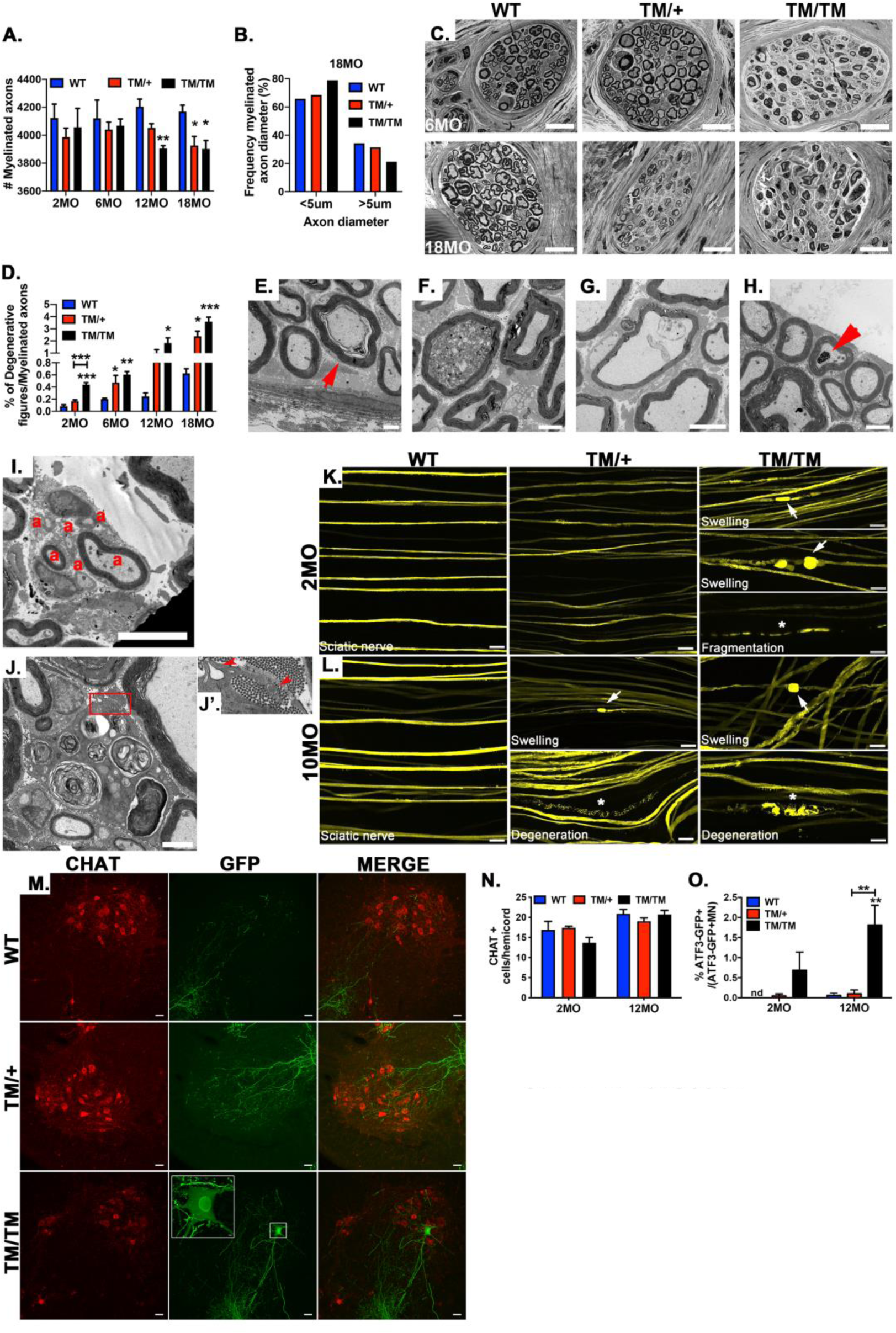
P0T124M mutation causes axonal degeneration. (**A**) Quantification of myelinated axons in whole sciatic nerves at 2, 6, 12, and 18 months of age. Starting at 12 months of age, the number of myelinated axons is significantly reduced in mice harboring the P0T124M mutation (TM mice). (**B**) Frequency distribution of myelinated axons as a function of axonal diameter shows that big axons (>5 μm in diameter) are slightly more impacted than small axons (<5 μm), especially in 18-month-old TM homozygous (TM/TM) mice. (**C**) Semithin cross sections of digital (toe) nerves show presence of degenerative fibers in 6-month-old TM/TM mice. At 18 months of age, TM/+ and TM/TM mice exhibit degenerative fibers. Scale bars: 20 μm. (**D**) Quantification of degenerative figures in whole sciatic nerves at 2, 6, 12, and 18 months of age. (**E to J**) Representative electron micrographs of degenerative axons observed in TM mice. (**E**) Typical axonal cuffing with large periaxonal collar. (**F**) Swelled axon containing dark vesicles and mitochondria. (**G**) Empty axonal swelling. (**H**) Myelinated axon containing glycogenosomes (arrowhead). (**I**) Axonal regenerative clusters. Note the presence of hypomyelinated or amyelinated axons. “a” indicate axons (**J**) Degenerative fiber associated with a macrophage. (**J’**) Inset magnified 1.5× shows macrophage protrusions. **E**, **F**, **H**, and **J** scale bars: 2 μm; **G** and **I** scale bars: 5 μm. (**K and L**) Representative confocal microscopy images of wild-type (WT) and TM/TM–Thy1-YFP sciatic nerves. (**K**) Note the presence of axonal swellings (arrows) and axonal degeneration (asterisks) in TM/TM at 2 months of age. (**L**) Axonal swellings (arrows) and axonal degeneration (asterisks) are observed in TM/+ and TM/TM mice at 10 months of age. Scale bars: 20 μm. (**M**) Representative confocal microscopy images of spinal cords sections (L3) from 12-month-old WT and TM/TM–ATF3-GFP mice stained for choline acetyltransferase (CHAT; motoneurons) (red) and ATF3-GFP (green). Scale bars: 40 μm. High-magnification inset shows motoneuron expressing GFP under ATF3 promoter control. (**N**) Quantification of motoneurons (CHAT-positive cells) in hemicords of 2- and 12-month-old mice. (**O**) Percentages of motoneurons expressing ATF3-GFP are increased in TM/TM mice at 2 and 12 months of age. *n* (animals) ≥ 3 per genotype. \**p < 0.05*, \*\**p < 0.01* by multiple-comparisons Tukey’s *post hoc* tests after one-way ANOVA. Graphs indicate means ± SEMs.

Although compact myelin ultrastructure was not markedly altered by the P0T124M mutation, we noticed, comparable to that in T124M-CMT2J patients^14^, occasional supernumerary SC processes and onion bulbs (**Fig. 2A, B, and I**), suggesting some active de-/remyelination in TM/TM mice. These features were rare (0.1-0.2% of myelinated fibers) and did not progress over time (**Fig. 2J**) as would be expected in pure demyelinating neuropathy models^24, 25^. Finally, in TM/TM mice at least 6 months of age, we detected myelin balloons (**Fig. 2K and supplementary Fig. 2D**), which are swellings thought to be formed by fluid influx inside the intraperiod line of the myelin sheath. Pressure created from fluid influx compresses the axon against one side of the balloon and could be responsible for axonal degeneration^26^.

Although we only noticed subtle changes in the myelin sheath *per se*, the most striking phenotype observed in TM nerves was related to fibers degeneration (arrowheads **Fig. 2A to D and supplementary Fig. 2**). A morphometric analysis revealed a significant reduction of myelinated fibers in sciatic nerves from TM mice at 12 and 18 months of age (**Fig. 3A**). At the latter timepoint, we noticed a slight shift toward smaller axon diameters in TM/TM mice, suggesting potential loss of large-diameter axons (**Fig. 3B**). Starting at 6 months of age for TM/TM mice and 18 months of age for TM/+ mice, we observed a massive loss of fibers within digital nerves (in the toes), suggesting a predominant distal dying back axonal damage (**Fig. 3C and supplementary Fig. 4**). Accordingly, degenerative features became apparent in TM sciatic nerves starting at 2 months of age and progressively increased over time affecting at 18 months, 2% and 3.5% of myelinated fibers in TM/TM and TM/+ mice, respectively (**Fig. 3D**). Beginning as early as 2 months of age in TM mice, we observed the classic hallmarks of axonal degeneration, such as the detachment of compressed axons from the inner myelin loop leaving behind large periaxonal collars (**Fig. 3E**). These are typical features of CMT1X neuropathy and could impair SC-to-axon communication^27, 28^. We also observed axonal swelling in nerves from TM/TM mice. Some of the swollen axons contained dense axoplasma with an accumulation of organelles, mostly mitochondria and dark vacuoles, suggesting defects in axonal transport, whereas other swellings showed empty vacuoles (**Fig. 3F and G**). In TM mice 6 months of age and older, macrophages were detected that appeared to be clearing out degenerated fibers (**Fig. 3J and J′**). We observed additional features of axonal degeneration, such as axonal regenerative clusters like those seen in samples from T124M-CMT2J patients and sciatic nerves from TM/TM and TM/+ mice at 12 and 18 months of age, respectively (arrows in **supplementary Fig. 2C and D** and **Fig. 3I**). Regenerating axons were either amyelinated or hypomyelinated. Furthermore, in the axolemma from TM nerves, we detected glycogenosomes (vacuoles containing glycogen) (**Fig. 3H** arrowhead). Glycogenosomes are naturally present in axons of aged WT rats^29, 30^ but are more common in neuropathies associated with diabetes^31^ and toxically induced neuropathy^32^.

**Figure 4:**
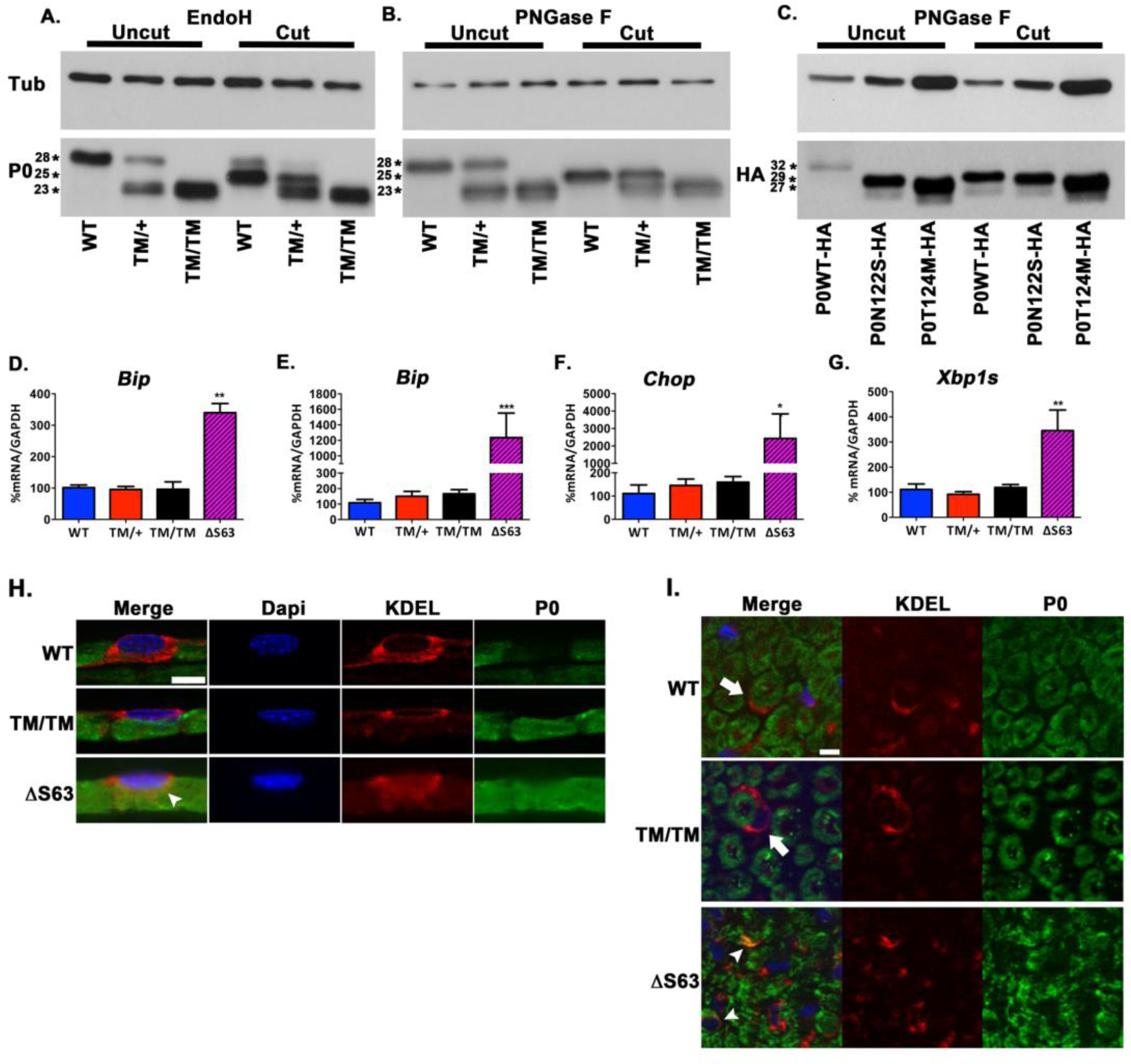
T124M mutation is responsible for molecular P0 modifications but does not alter P0 trafficking or UPR activation. Western blot analysis of P0 in sciatic nerve lysate from wild-type (WT), P0T124M heterozygous (TM/+), and homozygous (TM/TM) mice treated with (cut) or without (uncut) EndoH (**A**) and PNGase F (**B**). (**C**) COS-7 cells were transfected with P0-HA, P0T124M-HA, or P0N122S-HA. Samples were treated with (cut) or without (uncut) PNGase F and blotted with antibodies against HA. β-Tubulin (TUB) was used as a loading control. Asterisks indicate relative molecular weight. Bip, Chop, and Xbp1 spliced mRNAs were measured at postnatal day 10 (**D**) and postnatal day 30 (**E, F, and G**) by RT-qPCR TaqMan assay. GAPDH mRNA was used for normalization. ΔS63 sciatic nerve mRNA was used as a positive control. Representative immunofluorescence images of sciatic nerve teased fibers (**H**) and cross sections (**I**) at postnatal day 30 and 2 months of age, respectively. Fibers and cross sections were stained for P0 (green), endoplasmic reticulum (ER) (KDEL, red), and DAPI (blue). Note that P0T124M, like P0WT, is not retained in the ER and reaches the myelin sheath (arrows). P0ΔS63 is retained in the ER (arrowheads) and does not reach the myelin sheath. *n* (animals) ≥ 3 per genotype. \**p < 0.05*, \*\**p < 0.01*, \*\*\**p < 0.001* by multiple- comparisons Tukey’s *post hoc* tests after one-way ANOVA. Graphs indicate means ± SEMs.

To further assess axonal degeneration, we crossbred TM mice with two different transgenic reporter mouse lines. First, we crossed TM mice with Thy1-YFP reporter mice, which express yellow fluorescent protein in a small proportion of neurons^33^. In TM/TM mice as young as 2 months old, we identified bulb-like structures (swellings) at the tips of degenerating axons. They occasionally had a fragmented appearance typical of that seen during Wallerian degeneration after nerve transection or crush (**Fig. 3K**). At 10–11 months, TM/TM mice manifest dramatic axonal degeneration, and TM/+ mice also exhibit some axonal swelling and axonal degeneration (**Fig. 3L**). Second, we crossed TM mice with the ATF3-GFP reporter mouse line. Activating transcription factor 3 (ATF3) is a transcription factor rapidly expressed in neurons and SC after injury and axonal stress; intact adult neurons do not express ATF3^34^. The expression of ATF3 after sciatic nerve injury prevents cell death and promotes neurite formation and elongation, leading to enhanced nerve regeneration through transcriptional induction of survival and growth-associated genes^35^. As expected, in 2- and 12-month-old TM/TM–ATF3-GFP animals, ATF3-GFP was detected in motoneuron cell bodies (Choline acetyltransferase [ChAT] positive) from L1 to L5 spinal cord sections. However, in TM/+ mice, ATF3-GFP expression was marginal at 2 and 12 months of age (**Fig. 3M and O and supplementary Fig. 5**). Consistent with a peripheral neuropathy, the total number of motoneurons was not significantly changed (**Fig. 3N**).

**Figure 5:**
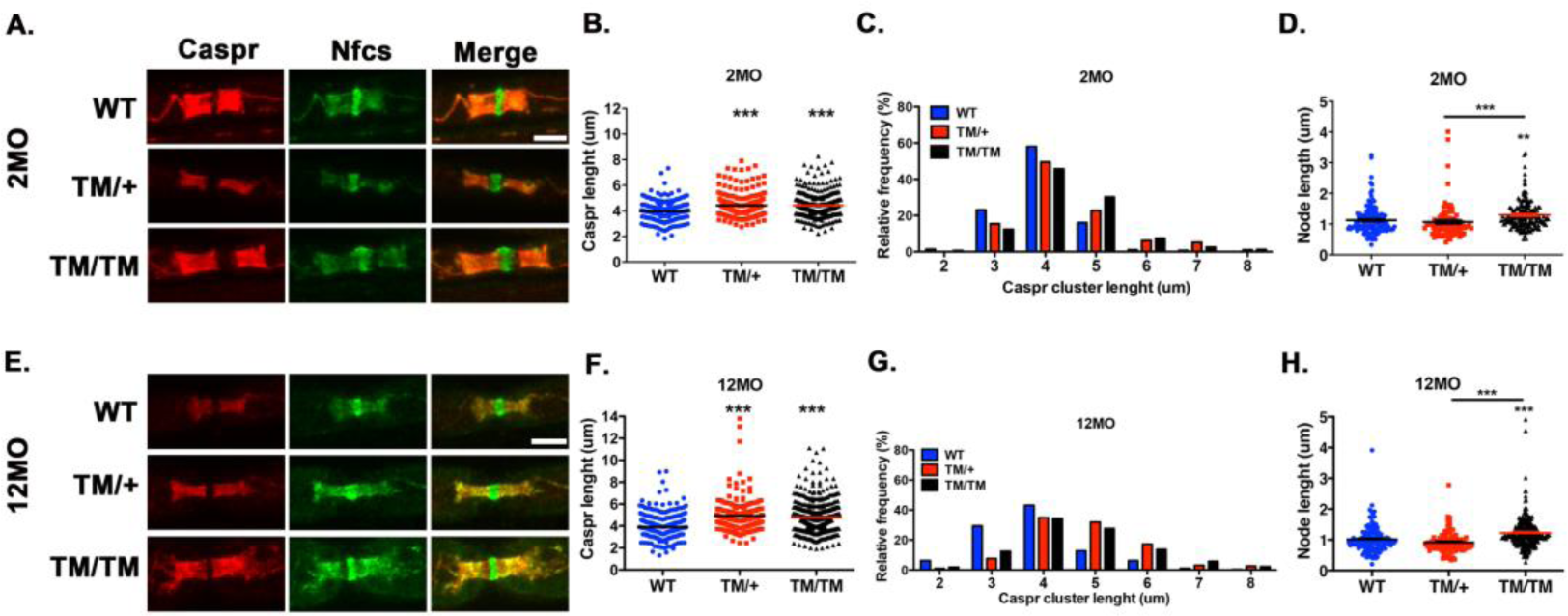
P0T124M mutation alters nodes and paranodes. Representative confocal pictures of sciatic nerve teased fibers stained with antibodies against the paranodal marker Caspr (red) and the paranodal and nodal marker pan-Neurofascin (Nfcs; green) at 2 (**A**) and 12 (**E**) months of age. Scale bars: 5 μm. Caspr length quantifications at 2 (**B**) and 12 (**F**) months of age. Relative frequency distributions of paranodal (Caspr) length at 2 (**C**) and 12 (**G**) months of age. Nodal length quantifications at 2 (**D**) and 12 (**H**) months of age. At least 200 paranodes and 100 nodes per genotype were quantified at each time point. *n* (animals) ≥ 3 per genotype. \*\*\**p < 0.001* by multiple-comparisons Tukey’s *post hoc* tests after one-way ANOVA. Graphs indicate means ± SEMs.

Altogether, our findings indicate that the TM mouse model closely replicates the clinical findings and axonopathy described in T124M-CMT2J patients. Even though P0 is expressed only by SC, we noticed only subtle changes in the TM compact myelin sheath, suggesting primary axon damage and Wallerian-like degeneration.

### P0T124M mutation impedes N-glycosylation

A denaturing Western blot revealed that P0T124M has a lower relative molecular weight than the WT protein (P0WT) (23 kDa and 28 kDa, respectively) (**supplementary Fig. 3 and Fig. 4. A and B**). The methionine residue in place of threonine 124, which is within the acceptor sequence for *N*-glycosylation, may impede P0 *N*-glycosylation, resulting in the observed shift in migration. To test this, we removed sugars of *N*-glycosylation via enzymatic digestion with endoglycosidase H (EndoH) or peptide N-glycosidase F (PNGaseF), which decreased the molecular weight of P0WT to 25 kDa. By contrast, P0T124M was insensitive to digestion by both enzymes, suggesting that P0T124M is not *N*-glycosylated. However, the lack of glycosylation is not sufficient to explain the observed shift in the molecular weight of P0T124M, as *N*-glycosylation accounts for only 3 kDa (**Fig. 4A and B**). To confirm this experimentally, we transfected an HA-tagged version of P0N122S, a non-glycosylatable mutant into COS-7 cells, and observed that it migrated at a higher relative molecular weight (29 kDa, comparable to deglycosylated P0WT-HA) than HA-tagged P0T124M (27 kDa) (**Fig. 4C**). Therefore, additional modifications, likely independent of *N*-glycosylation and specific to the T124M mutation, may be responsible for the altered migration of P0T124M.

### Pathogenesis of P0T124M does not involve the unfolded protein response

*N*-Glycosylation occurs in the endoplasmic reticulum and Golgi apparatus, and it is important for proper protein folding and for protein quality control^36^. The lack of *N*-glycosylation could impede P0T124M trafficking and generate endoplasmic reticulum stress. For example, the unfolded protein response is activated in two CMT1B mouse models: P0ΔS63^24, 37^ and P0R98C^17^. However, the levels of mRNA for unfolded protein response markers (binding immunoglobulin protein [BIP], C/EBP homologous protein [CHOP], and X-box binding protein 1 spliced [XBP1s]) were not increased in sciatic nerves from TM mice (**Fig. 4D to G**). Moreover, the trafficking of P0T124M was not altered, as the mutant protein was not retained in the endoplasmic reticulum and was able to reach the plasma membrane (**Fig. 4H and I**). These results suggest that the unfolded protein response is not involved in the pathogenesis of T124M-CMT2J disease.

### P0T124M mutation perturbs axon-glia interactions

Paranodes are specialized regions for axon-glia interaction that permit the diffusion of metabolites, hormones, and water-soluble molecules from SC to the periaxonal space^38^. Defects in paranodes are associated with axonal degeneration^39–41^, and P0 was shown to help maintain paranodal and nodal structures^42^. To detect potential defects in paranodal and nodal regions caused by the P0T124M mutation, they were stained with the markers Contactin associated protein 1 (Caspr) and pan-Neurofascin (Nfcs) in isolated teased fibers. At 2 months of age, the paranodal region in TM fibers was longer than in WT fibers (**Fig. 5A to C**). The frequency distribution of Caspr clusters was shifted toward increased lengths in TM mice relative to WT mice. Nodal length was also increased in TM/TM mice (**Fig. 5D**). Similar results were obtained from 12-month-old mice (**Fig. 5E to H**), confirming the elongation of paranodal and nodal areas in TM mice.

P0 is also important for the creation of Schmidt-Lanterman incisures (SLI), another non-compact myelin domain crucial for the transit of signals and molecules between the outer and inner SC surface and the axon^43^. Morphological analyses of TM sciatic nerves revealed a significant increase in SLI in a time- and gene dose-dependent manner (**Fig. 2B and C**, **supplementary Fig. 2** asterisks, and **Fig. 6A to D**). This was confirmed by staining for actin, Nfcs, and MAG, three proteins enriched in SLI, in isolated teased fibers from 2-, 6-, and 12- month-old animals (**Fig. 6E, I, M, and Q**). In mice as young as 2 months of age, the SLI in TM fibers were shorter (**Fig. 6F, J, N, and R**). Moreover, the morphology of SLI was also altered. Fibers from WT mice showed the expected series of single conical and clearly separated SLI, whereas the SLI in TM mice seem disorganized, more numerous, and sometimes fused together. By measuring the distance between two adjacent SLI, we confirmed that the number of SLI was increased in TM mice (**Fig. 6G, K, O, and S**). Finally, the number of SLI per 100 μm in TM/TM mice was 2-fold higher than in WT mice (**Fig. 6H, L, P, and T)**. These results suggest that P0T124M impairs the formation, structure, or function of SLI, which could ultimately result in lack of axonal metabolic support.

**Figure 6:**
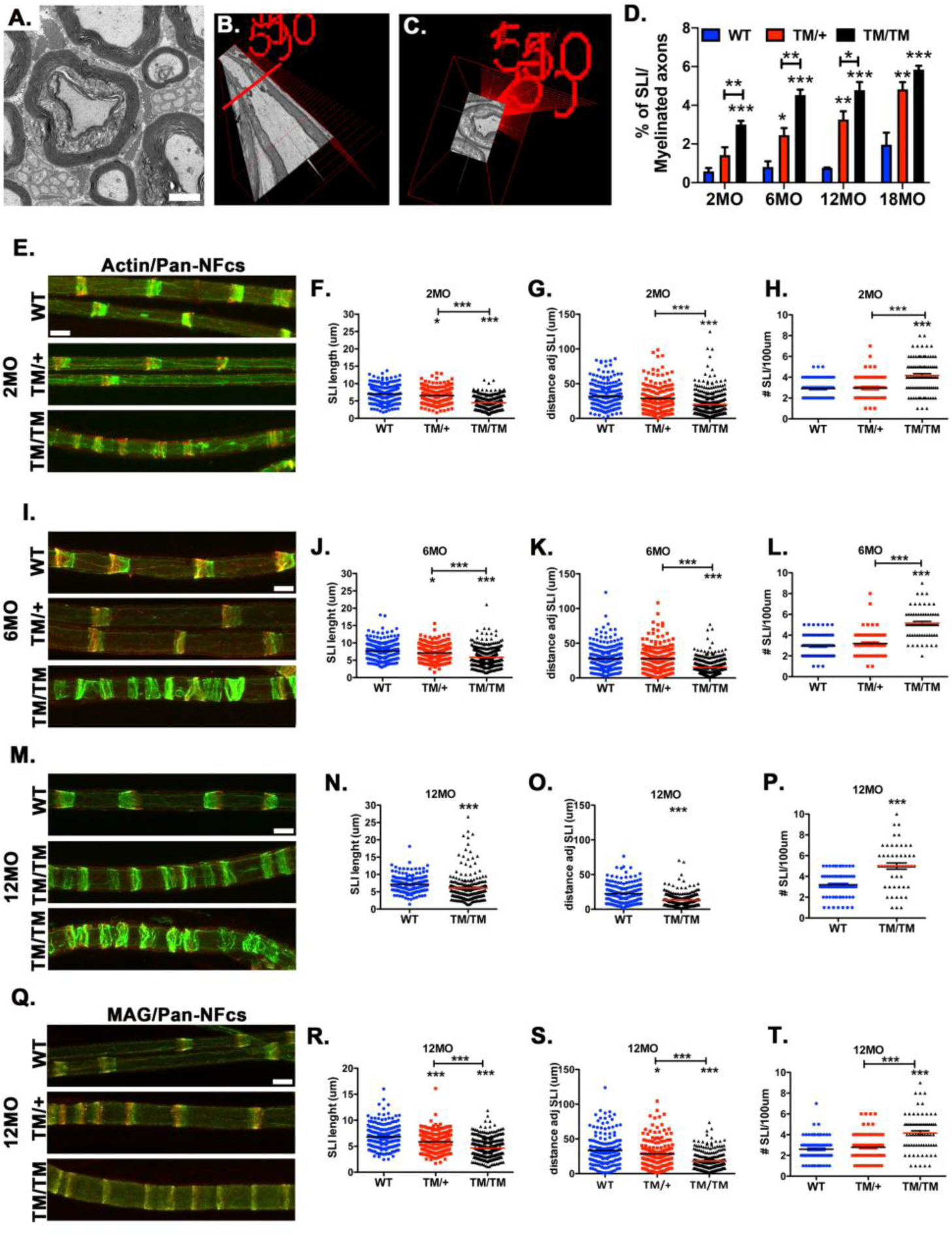
P0T124M mutation alters SLI morphology, length, and distribution. (**A**) Representative electron micrograph of Schmidt-Lanterman incisures (SLI) in sciatic nerve transverse section. SLI is illustrated by uncompacted myelin surrounded by two layers of compact myelin. (**B**) Three-dimensional electron micrograph showing myelinated axon with one SLI in longitudinal view. (**C**) Coronal view of the same SLI seen in B at the level of the red line. (**D**) Quantification shows an increased percentage of SLIs in nerves from mice harboring the P0T124M mutation (TM) at 2, 6, 12, and 18 months of age. Representative confocal pictures of sciatic teased fibers stained with FITC-phalloidin (actin, green) (**E, I, and M**) or anti-MAG antibodies (**Q**) (green) and anti-pan-Nfcs antibodies (red) at 2 (**E**), 6 (**I**), and 12 (**M and Q**) months of age. SLI morphology is disrupted in TM mice. Scale bar: 10 μm. Measurements of SLI length at 2 (**F**), 6 (**J**), and 12 (**N and R**) months of age. Measurements of the distance between adjacent SLI at 2 (**G**), 6 (**K**), and 12 (**O and S**) months of age. Quantifications of SLI number per 100 μm at 2 (**H**), 6 (**L**), and 12 (**P and T**) months of age. At least 207 SLI per genotype were quantified at each time point. *n* (animals) ≥ 3 per genotype. \**p < 0.05*, \*\**p < 0.01*, \*\*\**p < 0.001* by multiple-comparisons Tukey’s *post hoc* tests after one-way ANOVA (**D**, **F**, **G**, **H**, **J**, **K**, **L**, **R**, **S**, and **T**) or by two-tailed Student’s *t* test (**N**, **O**, and **P**). Graphs indicate means ± SEMs.

Connexin 32 (Cx32) is an important protein in the PNS. Similarly to P0T124M, Cx32 mutations cause CMT1X, characterized by axonal abnormalities that precede demyelination^44^. Cx32 is located in non-compact myelin domains (SLI and paranodes), where it forms gap junctions between the layers of the SC myelin sheath, providing a fast diffusive radial path between the abaxonal and adaxonal areas^45^. Although we were not able to reproducibly detect Cx32 in SLI, we quantified the fluorescence intensity of Cx32 staining in paranodes of WT and TM mice at 2 months of age (**Fig. 7A**). Compared to WT mice, the intensity of Cx32 staining was 40% lower in nerves from TM/TM mice but similar to or even slightly higher in TM/+ mice (**Fig. 7B and E**). Cx32 was also mislocalized in some TM fibers. While 90% of paranodal Cx32 colocalized with Caspr in WT fibers, 20% of TM/+ fibers and 30% of TM/TM fibers showed Cx32 outside the paranodal area (**Fig. 7C**). To confirm Cx32 mislocation in TM mutants, we costained teased fibers for Cx32 and potassium voltage-gated channel 1.1 (KV1.1), a marker of the juxtaparanodal region (**Fig. 7D**). Cx32 was expressed in juxtaparanodes (i.e., colocalized with KV1.1) in only 15% of WT fibers but in 20% and 35% of fibers from TM/+ and TM/TM mice, respectively (**Fig. 7F**). It is tempting to speculate that the altered Cx32 distribution contributes to the axonopathy observed in TM mice.

**Figure 7:**
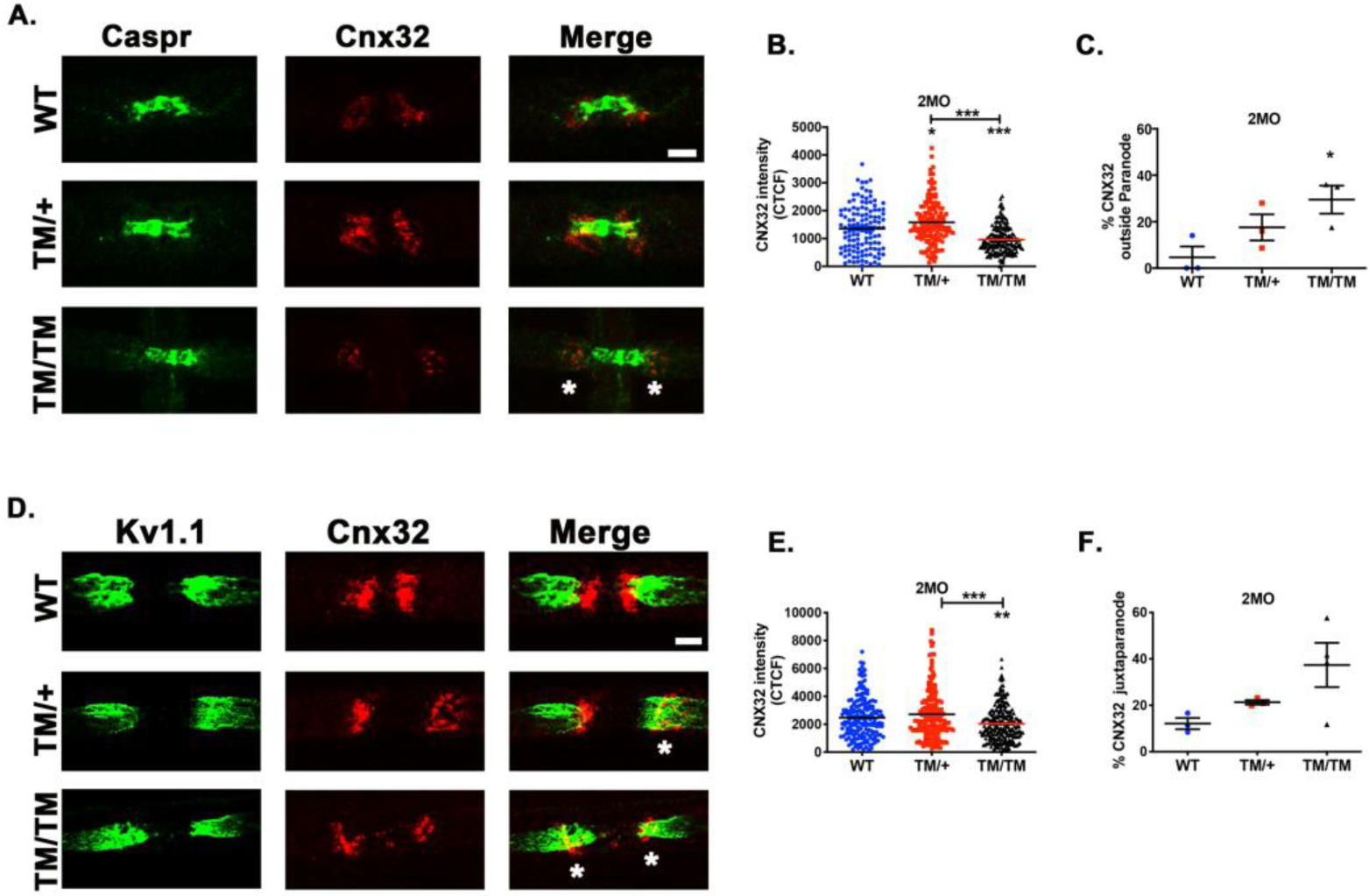
P0T124M results in altered Cnx32 expression. Representative confocal pictures of sciatic nerve teased fibers stained with antibodies against connexin 32 (Cnx32) (red) and Caspr (green) (**A**) or Kv1.1 (green) (**D**) from mice at 2 months of age. Asterisks indicate Cnx32 signal located at juxtaparanode. Scale bars: 5 μm. (**B and E**) Quantifications of the intensity of Cnx32 staining (measured by corrected total cell fluorescence methodology). Quantifications of Cnx32 signals expressed at juxtaparanodes: not colocalized with Caspr (**C**) and colocalized with Kv1.1 (**F**). Cnx32 intensity was quantified from at least 122 para- and juxtaparanodes per genotype. *n* (animals) ≥ 3 per genotype. \**p < 0.05*, \*\**p < 0.01*, \*\*\**p < 0.001* by multiple- comparisons Tukey’s *post hoc* tests after one-way ANOVA. Graphs indicate means ± SEMs.

Altogether, our results indicate a perturbation of the areas of non-compact myelin where exchange and communication between SC and axons are thought to occur. These alterations could lead to deficient communication and support from SC to axons.

### ATP and NAD^+^ but not glycolysis are decreased in T124M mice

Because of the alterations at axon-glia exchange areas, we reasoned that metabolite transport from SC to axons would be disturbed in TM mice. We studied the steady state levels of key metabolites for SC and axon functions by mass spectrometry in sciatic nerves from 12-month-old mice (**Fig. 8A to H**). ATP, the main energy source for neurons, decreased by 30% in TM/TM mice (**Fig. 8A**). Glycolytic activity in myelinating glia is fundamental for axonal energetics and survival^46–51^. However, there were no significant differences between WT and TM/TM mice in the amounts of glucose-6-phosphate (G6P) (**Fig. 8B**), lactate (**Fig. 8C**), or pyruvate (**Fig. 8D**). Interestingly, there was striking and significant reduction (-55%) in the amount of oxidized nicotinamide adenine dinucleotide (NAD^+^) in sciatic nerves from TM/TM mice (**Fig. 8E**) along with a trend toward less reduced NAD (NADH) (**Fig. 8F**). NAD^+^ and NADH are involved in many cellular and biological functions such as energy metabolism, mitochondrial function, and redox state^52^, each of which is important for axonal physiology. Moreover, NAD^+^ balance is crucial for axonal survival. Cleavage of NAD^+^ by SARM1 (sterile alpha and toll/interleukin 1 receptor motif-containing 1) into nicotinamide (NAM) and adenosine phosphoribose (ADPR) leads irremediably to axonal degeneration^53–55^. Although ADPR expression appears to be normal in TM nerves (**Fig. 8G**), it is rapidly converted to cyclic ADPR (cADPR), making it difficult to obtain a reliable measurement. However, NAM expression shows a trend toward increased levels in TM/TM mice (**Fig. 8H**). This combined with the reduced levels of NAD^+^ suggest an involvement of SARM1 in the axonal degeneration observed in TM mice.

**Figure 8:**
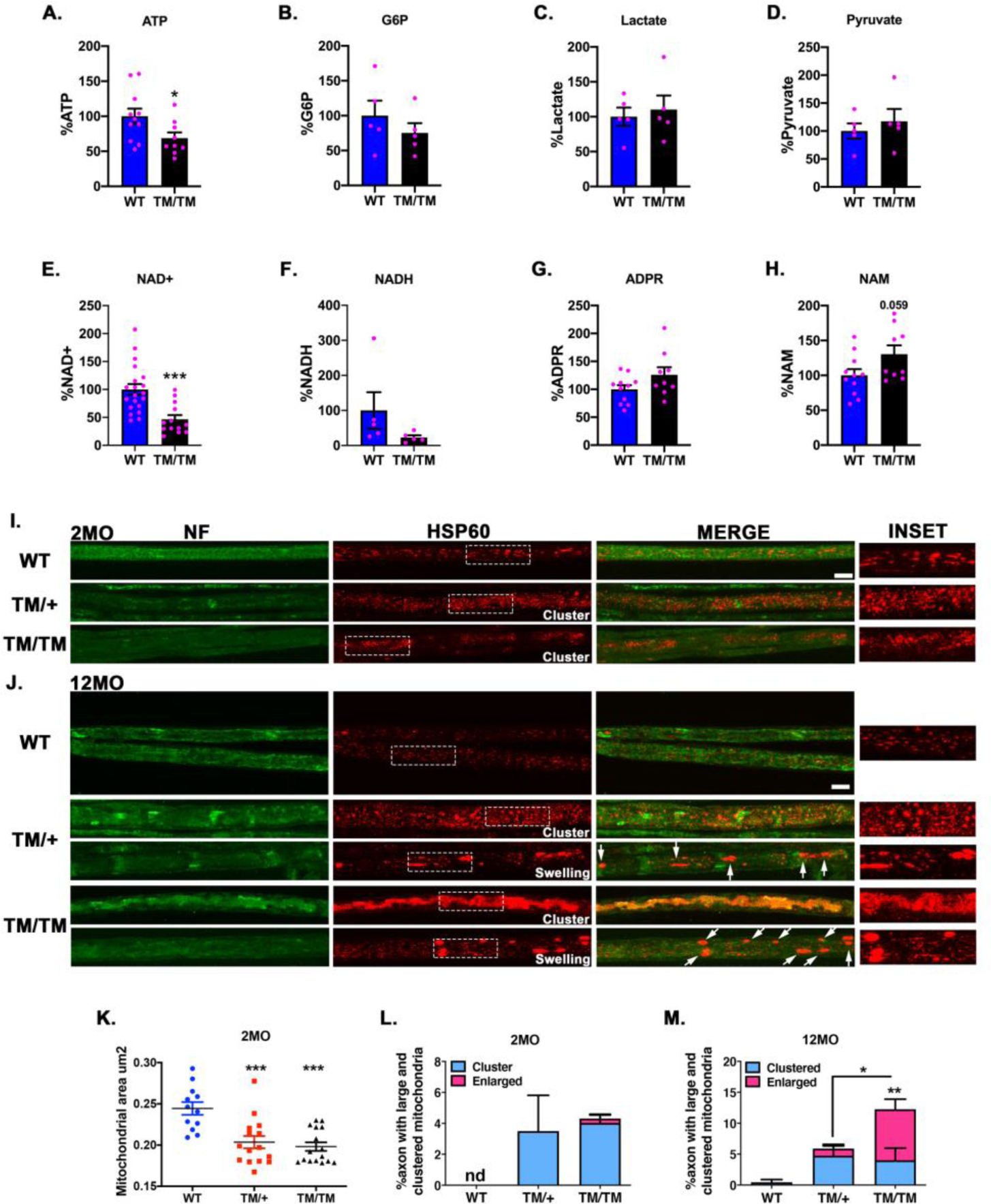
Metabolic and axonal mitochondrial impairments in TM mice. (**A to H**) Measurements of metabolites involved in axonal energy and degeneration programs. ATP (**A**), G6P (**B**), lactate (**C**), pyruvate (**D**), NAD^+^ (**E**), NADH (**F**), ADPR (**G**), and NAM (**H**) were quantified via liquid chromatography-tandem mass spectrometry of sciatic nerves from 12- month-old mice. *n* (animals) ≥ 5 per genotype. (**I and J**) Representative confocal pictures of sciatic nerve teased fibers stained with antibodies against mitochondrial protein HSP60 (red) and axonal marker neurofilament (NF) (green) from mice at 2 (**I**) and 12 (**J**) months of age. Arrows indicate swelled mitochondria. Scale bars: 10 μm. High magnification insets show mitochondria. (**K**) Quantification of mitochondrial surface area. Surface area was quantified from at least 1,096 mitochondria per animal. Proportions of axons with clustered and enlarged mitochondria at 2 (**L**) and 12 (**M**) months of age. At least 263 axons per genotype were imaged. *n* (animals) ≥ 3 per genotype. \**p < 0.05*, \*\**p < 0.01*, \*\*\**p < 0.001* by two-tailed Student’s *t* test (**A** to **H**) and by multiple-comparisons Tukey’s *post hoc* tests after one-way ANOVA (**K** to **M**). Graphs indicate means ± SEMs.

### Axonal transport and mitochondrial disruption in T124M mice

Because ATP levels were decreased in TM mice and because neurons are dependent on mitochondria for ATP homeostasis and long-term integrity, we posited that the TM mice would exhibit axonal mitochondrial defects. We stained mitochondria in sciatic nerve teased fibers with anti-HSP60 (heat shock protein 60) antibodies (**Fig. 8I and J**). By co-staining with neurofilaments and using confocal microscopy, we focused our attention to axonal mitochondria. In TM fibers, axonal mitochondria appeared fragmented. We observed a global decrease of axonal mitochondrial surface area in TM mutants compared to that in WT (**Fig. 8K**). We also noticed, starting at 2 months of age, the presence of clustered axonal mitochondria in approximately 4 to 5% of TM axons, suggesting a defect in fast axonal transport **(Fig.8L)**. Moreover, at 12 months of age, large axonal mitochondria were present in 1 and 10% of TM/+ and TM/TM fibers respectively (**Fig.8J, arrows and 8M**). Large mitochondria could reflect clusters of small mitochondria or degenerative swelling. Altogether, our results indicate a defect in the transport and, potentially, functionality of axonal mitochondria in sciatic nerves in TM mice that in turn could impair ATP production, thereby leading to axonal death.

## Discussion

We generated a genetically authentic knock-in mouse model of the human inherited neuropathy CMT2J caused by the T124M mutation encoded in the *MPZ* gene. In-depth characterization of these mice demonstrated that mutants closely replicate the axonopathy and other clinical aspects of T124M-CMT2J patients: an adult-onset progressive motor and auditory neuropathy with a marked reduction of CMAP, but only a slight decrease of NCV at later stages; reduced numbers of myelinated axons and the presence of regenerative clusters, but also well compacted myelin sheaths with near normal packing and periodicity with only occasional onion bulbs and supernumerary SC^14, 56, 57^. Although the presence of onion bulbs may suggest sporadic demyelination in TM/TM mice, our findings, together with the clinical features of the disease and the fact that changes in axonal changes preceded those in compact myelin, suggest that demyelination is not likely to trigger axonal degeneration in these mice.

Despite the similarities, we also identified differences between TM mice and patients. The mouse axonal neuropathy appears to be milder than its human counterpart. Humans present with an axonal loss of 50–70%^14, 15^, whereas the loss is around 10% in mice. The allelic pattern is also different between mice and humans. T124M-CMT2J is an autosomal dominant disease. All CMT2J patients referenced were heterozygous for the P0T124M mutation except for one patient that was homozygous for the mutation; this patient was severely affected and died at the age of 44 years^56, 57^. Although TM/+ mice similarly develop the disease, the human disease phenotype was more faithfully recapitulated in TM/TM mice. The milder severity and allelic pattern discrepancy are common observations in CMT2 knock-in mouse models^58^ and could be explained by several factors, including the shorter life span and shorter axonal length in mice.

Another difference between humans and mice is myelin sheath thickness. In patients, the thickness of sural nerve myelin is varied. In general, human myelin sheaths are thin, but some patients have normal or even thickened myelin sheaths^14, 15^. In TM mice, we did not observe hypomyelination. Myelin sheath thickness was normal until 6 months of age, and at 12 months of age, TM fibers were slightly hypermyelinated.

Homophilic interactions of P0 proteins are fundamental for myelin compaction. P0 interacts with itself via the protein backbone, and P0 carbohydrates may contribute to homophilic adhesion^59, 60^. Even though the P0T124M mutation abolished P0 *N*-glycosylation, the myelin sheath was relatively well compacted. Moreover, *in vitro* experiments have shown that the T124M mutation does not interfere with P0 adhesiveness^61^. Interestingly, the P0T124A and P0N122S mutations, which similarly abolish native *N*-glycosylation, are responsible for late-onset CMT1B with only moderate reduction of NCV^62, 63^, and like P0T124M, these mutated P0 proteins retain their adhesive propriety *in vitro* (Dr. Francesca Veneri personal communications). This suggests that the P0 protein backbone is generally sufficient for P0 homophilic adhesion and for myelin compaction. On the other hand, we cannot exclude the possibility that the absence of *N*-glycan has subtle consequences for P0 adhesion and myelin stability, which may explain the observed onion bulbs and hypermyelination. Future analyses of TM myelin periodicity by X-ray diffraction would answer this question.

Notably, the P0T124M mutation altered non-compact myelin, which comprises domains essential for SC-to-axon interactions. Indeed, in TM fibers, SLI were shorter, more numerous, and had a disorganized structure. The P0T124M mutation also altered paranodal structures. The aberrant Caspr immunoreactivity we observed could disrupt paranodal–axoglial interactions^39^. Moreover, the dialogue between SC and axons is likely impaired in fibers with large periaxonal collars, as the adaxonal SC membrane cannot contact the underlying axons. Altogether, our results strongly suggest that axonal degeneration in TM mice is caused by the disruption of the physical and molecular communication between glia and axons.

How P0 is involved in neuroglial interactions and signaling at SLI and paranodes is an important but unanswered question. P0 is not a structural component of SLI, but it appears to be necessary for SLI formation via an unknown mechanism^43, 64^. Moreover P0, via interaction with Nfcs, was shown to be directly involved in paranode maintenance^42^. Future experiments are needed to determine if the P0T124M mutation disrupts this interaction. It is tempting to speculate that the lack of P0 *N*-glycans as a result of the T124M mutation is detrimental for SC-to-axon interactions. Indeed, *N*-glycans modifications (GlcNAc-6-*O*-sulfation) are directly involved in the maintenance of paranodal organization in the PNS^41^.

Because of the defect in SC-to-axon exchange areas and because myelinating glia and axons are metabolically linked^49, 65, 66^, we suspected that the metabolic support from SC to axons is interrupted in TM mice. This hypothesis is further reinforced by the reduced ATP levels observed in nerves from TM mice. In the CNS, oligodendrocyte glycolysis through lactate production is indispensable for producing the ATP needed for axonal activity and survival^46, 47, 50, 51^. However, despite the reduced ATP and NAD^+^ levels, which are crucial for glycolysis and oxidative phosphorylation^53, 67^, glucose uptake (G6P) and glycolysis (lactate and pyruvate) were not altered in TM nerves. In the PNS, SC shift towards glycolysis in favor of lactate production to sustain axons after acute injury^68^. However, the lactate and glycolysis pathways play a limited role in axon survival in the PNS under physiological conditions^48, 68–70^. Other metabolic pathways, such as lipid metabolism, may be more important for axonal health in the PNS^48^. Thus, TM mice described here represent a highly valuable model to further our understanding of SC-to-axon metabolic interactions, and the identification of altered metabolic pathways could pinpoint potential therapeutic targets^71–74^.

The mitochondrial defects observed further support the notion that P0T124M disrupts the metabolic support of axons. Neurons depend on mitochondrial ATP production to survive and are, therefore, particularly vulnerable to changes in mitochondrial morphology and connectivity. Our results suggest that perturbation of SC-to-axon support that results from the P0T124M mutation has deleterious consequences for the morphology of axonal mitochondria, illustrated by a global reduction of axonal mitochondrial surface (fragmentation) and by the presence of clustered and swollen mitochondria. The relationship between mitochondrial fragmentation and bioenergetics is bidirectional because mitochondrial fragmentation decreases ATP production, which in turn induces mitochondrial fragmentation^75^. The presence of clustered and swollen mitochondria in the nerves from TM mice also hints to a potential impairment in fast axonal transport, to which axons, because of their considerable length, are particularly sensitive^76^. A disruption of energy-dependent axonal transport is compatible with the distal dying back mechanism of axonal degeneration observed in TM mice. Furthermore, axonal transport defects and clustered and swollen mitochondria are hallmarks of degenerating axons in myelin proteolipid lipid (PLP)-null mice^77^ and P0-CNS mice, a transgenic mouse in which PLP expression was artificially substituted by P0 in oligodendrocyte myelin^78^.

Mitochondrial dysfunction^55, 79^ and decreased levels of NAD^+^ coupled to the trend of increased NAM levels^54^ also point to a role for SARM1 in the axonal degeneration observed in TM mice. SARM1 is considered a very interesting therapeutic target to limit axonal degeneration in many neurodegenerative diseases, and such approaches may also apply to T124M-CMT2J.

## Concluding remarks

The authentic mouse model for T124M-CMT2J we generated shows that the P0T124M mutation alters paranode, SLI, and SC gap junction structures, possibly resulting in a defect of trophic support from SC to axons and energetic failure. We suspect that the initial ATP deficiency results in axonal mitochondrial dysfunction, which in turn worsens the axonopathy. Axonal transport is slowed, leading to a vicious cycle that ultimately results in axonal degeneration with potential involvement of the SARM1 cascade (**supplementary Fig. 6**). The fidelity of the TM mouse model to CMT2J disease observed in patients and the understanding of the pathomechanisms underlying SC–axon communication defects will enable us to explore new therapeutic strategies for T124M-CMT2J disease as well as other CMT and neurological diseases that involve axonal degeneration.

## Supporting information

Detailed Materials and Methods

sequencing

## Acknowledgements

We thank Courtney Williamson and Anke Claessens for their excellent technical assistance, Dr. Karen Dietz for assistance preparing this manuscript.

## Funding

We are grateful for funding from: Charcot-Marie-Tooth association (to L.W and G.G.S) and Fondazione Telethon (GGP19099 to M.D)

## Competing interests

The authors report no competing interests.

## Supplementary material

Supplementary material is available at *Brain* online.

## Notes

### Competing Interest Statement

The authors have declared no competing interest.

