## Supplementary material for "MPZ-T124M mouse model replicates human axonopathy and suggest alteration in axo-glia communication": Detailed Materials and Methods

### Animals

The P0T124M line was established in two different backgrounds: C57BL/6N and FVB/N. All results were obtained using the P0T124M C57BL/6N line, except for those from auditory system (**Fig. 1L, J and K**) and morphology (**supplementary Fig. 4)** experiments, in which the P0T124M FVB/N line was used. Progeny used in this study were N3–N10 congenics in either the C57BL/6N or FVB/N background. For the detailed characterization of axonal morphology and neuronal stress, P0T124M mice were crossed with B6.Cg-Tg(Thy1-YFP)HJrs/J mice (JAX# 003709), here called Thy1-YFP, and ATF3-GFP mice. ATF3-GFP mice were kindly provided by Dr. Clifford J. Woolf (Harvard Medical School).

Animals were housed no more than five animals per cage with a 12-h light/dark cycle. Mutant and control littermates from either sex were sacrificed at the ages indicated in the text.

### Genotyping and PCR primers

Genotyping was performed by PCR analysis of genomic DNA extracted from toe clips or ear punches. In addition to the NSP1 strategy described in the text, a loxP site was also used to genotype P0T124M mutants. PCR primers MpzInt3 Fw (5′-TCAAAGAGGGTGTCAGGGAG-3′) and MpzInt3 Rv (5′-GTGGCCCAGATTGGTCTTTA-3′) generate a 355- or 305-bp amplimer for loxP or wild type, respectively. For ABR and cochleogram experiments, cadherin 23 (*cdh23*) polymorphisms were genotyped. PCR primers Cdh23 Fw (5′-ATCATCACGGACATGCAAGA-3′) and Cdh23 Rv (5′-AGCTACCAGGAACAGCTTGG-3′) generate a 315-bp amplimer. After digestion with BSRI, the C57BL/6N allele migrates at 238 bp + 77 bp, whereas the FVB/N allele is not sensitive to BSRI digestion. Instead, after BSAWI digestion, the FVB/N allele migrates at 236 bp + 79 bp, whereas the C57BL/6N allele is not sensitive to BSAWI digestion. Only animals with at least one *cdh23* FVB/N allele were used for experiments. Thy1-YFP mouse genotyping was performed as described by Jackson Laboratory (https://www.jax.org/strain/003782). PCR primers Thy-YFP Fw (5′-ACAGACACACACCCAGGACA-3′) and Thy-YFP Rv (5′-CGGTGGTGCAGATGAACTT-3′) generate a 400-bp amplimer. PCR primers for internal control

Fw (5′-CTAGGCCACAGAATTGAAAGATCT-3′) and internal control Rv (5′-GTAGGTGGAAATTCTAGCATCATCC-3′) generate a 300-bp amplimer. PCR primers ATF3-GFP Fw (5′-CAACATCCTGTTCGGCAACCAA-3′) and ATF3-GFP Rv (5′-TCCACGCGGTACACGAACTT-3′) generate a 244-bp amplimer.

### Isolation of genomic DNA for genotyping

Toes and ear punches were digested in 75 µL of 25 mM NaOH-0.2 mM EDTA at 95°C for 45 min and neutralized with 75 µL of 40 mM Tris-HCl (pH 5.5). Samples were centrifuged and used for PCR reactions.

### Cell culture and transfection

COS-7 cells were obtained from the American Type Tissue Collection. Cells were grown in low-glucose Dulbecco's modified Eagle's medium supplemented with 10% fetal bovine serum and antibiotics (100 µg/ml penicillin-streptomycin) in a humidified atmosphere containing 5% $CO_2$ at 37°C. For transient transfection, Lipofectamine 2000 (Invitrogen) and 4 µg of plasmid DNA were incubated separately in Opti-MEM (Gibco-BRL) for 5 min at room temperature and then combined for another 20 min. When cells reached approximately 80% confluence, they were washed with Opti-MEM and then incubated with the combined Lipofectamine-DNA solution for 6 h at 37°C. The cells were then washed once with HBSS (free of calcium and magnesium) and incubated for 3 days in medium at 37°C before being processed for Western blotting.

### DNA constructs

HA-tagged P0WT and P0N122S plasmids used in these experiments were previously generated by Dr. Elisa Tinelli as described by Penutto *et al*[37]. HA-tagged P0T124M was generated by site-directed mutagenesis using a QuikChange II site-directed mutagenesis kit (Agilent Cat# 200523) according to the manufacturer's instructions.

P0WT-HA plasmid was used as a template. The following primers were used: Fw 5'-GGTTTTTGACATCACATGTGAACATGCCGTTGTCACTGTAGTCTAG-3'; Rv 5'-CTAGACTACAGTGACAACGGCATGTTCACATGTGATGTCAAAAACC-3'.

The P0T124M-HA sequence was confirmed by automated sequencing analysis.

### Behavioral tests

Accelerating rotarod analysis: Only female N3 C57BL/6N congenics at the ages indicated in the text were used. All mice were tested in a session of three trials per day for five consecutive days. Rotarod conditions were set to an acceleration of 5 rotation per minute$^2$ (rpm$^2$), starting at a minimum velocity of 4 rpm and accelerating to a maximum velocity of 40 rpm. Each trial consisted of one acclimating run that was not scored. The next three runs were recorded and averaged. Each run was stopped when the mouse fell or passed completely underneath the rod (180° of rotation) consecutively twice. Motor performance were determined from 5 mice per genotype at 2 months of age; 8 WT, 11 TM/+ and 9 TM/TM at 6 months of cage; 4 WT, 10 TM/+ and 7 TM/TM at 12 months of ages.

Beam walking test analysis: Both male and female N5 C57BL/6N congenics at 6 months of age were used. The beam apparatus (OpenScience Russia, Cat# TS0806-M) consists of 80-cm long beams with a flat surface 12- or 6-mm wide resting 50 cm above the tabletop on two poles with a slope of 15%. A black box is placed at the end of the beam as a finish point. A lamp, with a 60-watt light bulb, shines light above the start point and serves as an aversive stimulus. Thirty minutes before training/testing, the mice were transported to the room containing the beam apparatus. On training days (2 days), each mouse crossed the 12-mm-wide beam three times and then the 6-mm-wide beam three times (10-min rest between training sessions on the two beams). On the testing day, each mouse crossed the 6-mm beam four times, and each trial was video recorded. The time to cross the beam and the number of slips were measured and averaged. 6 mice per genotype were analyzed.

### Electrophysiological analysis

Electrophysiological analyses were performed as described by Weinstock *et al*[80]. Mice were anesthetized, at the ages indicated in the text, with 2,2,2 tribromoethanol (Avertin; Sigma-Aldrich), 20 mg/mL in $H_2O$, and placed under a heating lamp. The sciatic nerve conduction velocity was obtained with steel monopolar needle electrodes. One pair of stimulating electrodes was inserted subcutaneously near the nerve at the ankle, a second pair of electrodes was placed at the sciatic notch, and a third pair was placed over the dorsum of the spine. The compound motor action potential was recorded with an active electrode inserted into the muscles in the middle of the paw and a reference needle in the skin between the first and second digits. Electrophysiological parameters were determined from: 8 WT, 8 TM/+ and 6 TM/TM sciatic nerves (11 mice total) at 2 months of age, 14 WT, 18TM/+ and 20 TM/TM sciatic nerves (total 26 mice) at 6 months of age and 8WT, 12TM/+ and 14TM/TM sciatic nerves (total 17 mice) at 12 months of age.

### Simoa plasma Nfl measurment

Mice, at the ages indicated in the text, were anesthetized with 2,2,2 tribromoethanol (Avertin; Sigma-Aldrich), 20 mg/mL in $H_2O$. Five hundred microliters of blood per mouse was taken by cardiac puncture and placed in tubed coated with EDTA. Four microliters of EDTA was added to each sample and gently mixed. Samples were incubated for 15 min on ice and centrifugated for 10 min at 3,500 rpm at 4°C. Supernatants (plasma) were transferred to new tubes and stored at –80°C until analysis. Plasma sample Nfl concentration was determined using the in-house Simoa Nfl assay as described by Rohrer JD *et al*[81]. 9 mice per genotype were used at 2 months of age; 6 mice per genotype at 12 months of age.

### Morphological assessments

Mice at the ages indicated in the text were euthanized, and sciatic nerves and toes (digital nerves) were dissected, fixed in 2% glutaraldehyde, and stored at 4°C until processing. Nerves were then washed in phosphate buffer (0.12 M, pH 7.4), postfixed in 1% osmium tetroxide, dehydrated in increasing ethanol concentrations (50%, 70%, 90%, and 100%), incubated in propylene oxide, and finally embedded in Epon 100%. Nerves were then cut into semithin sections of 1-μm thickness or ultrathin sections of 80–85-nm thickness. Semithin sections were stained with toluidine blue 2% (phosphate buffer [0.12 M pH 7.4]). Ultrathin sections were stained with uranyl acetate (in

$dH_2O$) and lead citrate. For semithin sections, images were acquired with the 100× lens objective and stitched using PTGui software v.10 (New House Internet Services BV) to reconstruct a complete image of the nerve. Morphological parameters (degenerative figures, SLI and onion bulbs) were then evaluated for the full nerve. Electron micrographs were used for quantifications of g-ratios (at least 100 myelinated axons per animal), axonal size (at least 300 myelinated axons per animal), and myelin periodicity (10 myelin sheaths per animal) as described by Belin *et al*[82]. Schwann cells were identified by the presence of a basal lamina, whereas macrophages were identified by characteristic morphologic features, including microvilli. Quantifications were performed using ImageJ Fiji v1.52p. Between 3 and 6 mice per genotype per time point were analyzed.

For 3D EM reconstruction, tissue preparation and imaging were performed as previously described by Yin Xi *et al*[78]. In brief, mice at 12 months of age were perfused with 0.1-M sodium cacodylate buffer containing 2.5% glutaraldehyde (Electron Microscopy Sciences) and 4% paraformaldehyde. Sciatic nerves were dissected, cut into 5-mm segments, postfixed in 0.1% tannic acid in buffer and then stained successively with osmium ferricyanide, tetracarbohydrazide, aqueous osmium tetroxide, saturated aqueous uranyl acetate, and Walton's lead aspartate stains. Tissues were then dehydrated in graded ethanols and embedded in embedding resin at 60°C for 48 h. Nerve segments were mounted on aluminum pins, trimmed, surrounded with silver paste, and examined in a Sigma VP scanning electron microscope (ZEISS) fitted with a 3View in-chamber ultramicrotome (Gatan) and a low-kilovolt backscattered electron detector (Gatan). Longitudinally oriented axons were imaged covering areas ~80 µm wide and 80–400 µm in length. Images were collected at 1.8–2.5 kV, depending on tissue contrast, and up to 500 slices were cut at a thickness of 40–100 nm. Imaging was conducted using a 30-µm aperture in high-current mode and imaged at room temperature (21°C at a chamber vacuum of $10^{-6}$ mbar and working distance of 5.7 mm). Images were reconstructed and registered using ImageJ/FIJI software (National Institutes of Health).

### Trans-cardiac perfusion

Mice were anesthetized with 20 mg/mL 2,2,2 tribromoethanol (Avertin; Sigma-Aldrich) (0.02 mL/g of body weight). Once the mouse was unconscious, the thoracic wall was removed and the right atria was punctured. Twenty-five milliliters of 1× PBS was perfused into the left ventricle, followed immediately by 25 mL of 4% paraformaldehyde. The spinal cord was dissected and postfixed in 4% paraformaldehyde for 24 h at 4°C, followed by sucrose and OCT embedding. Tissues were frozen in OCT and stored at –80°C.

### Teased fiber preparation

Slides were coated with 3-aminopropyl-triethoxysilane (TESPA; Sigma-Aldrich) by subsequently submerging glass slides in acetone for 1 min, 4% TESPA in acetone for 2 min, and two times in 4% TESPA in acetone for 30 s each. Nerves were fixed in 4% paraformaldehyde for 30 min and then washed with 1× PBS. Nerves were desheathed with forceps and a 27-gauge needle. Bundles of fibers were separated in 1× PBS using 27-gauge needles and then gently teased apart to single fibers on TESPA-coated slides using modified insulin syringes containing minutien pins (Fine Science Tools #26002-10) attached to their needles. Slides were allowed to dry for at least 1 h and stored at –80°C. Sciatic nerves from 3 to 5 mice per genotype per time point were teased and analyzed.

### Immunofluorescence

Teased fibers and spinal cord (20 μm), and sciatic nerve (10 μm) cross sections were rehydrated in 1× PBS for 1 min and permeabilized with cold methanol. Tissues were rinsed in 1× PBS three times for 5 min and blocked with 5% fish skin gelatin and 0.5% Triton X-100 (teased fibers) or with 20% fetal bovine serum, 2% bovine serum albumin (BSA), and 0.1% Triton X-100 (cross sections) for 1 h at room temperature. The following primary antibodies were diluted in appropriate blocking buffer for incubation overnight: 1:300 chicken anti-P0 (Aves Lab Cat# PZO), 1:200 mouse anti-KDEL (Enzo Life Sciences Cat# ADI-SPA-827), 1:1,000 rabbit anti-Caspr (a gift from Dr. Elior Peles[83]), 1:500 chicken anti-pan-Neurofascin (R&D Systems Cat #AF3235), 1:200 rabbit anti-Kv1.1 (Alomone Cat# APC-009), 1:500 rabbit anti-HSPD1 (also known as HSP60) (Proteintech Cat# 15282-1-AP), 1:500 chicken anti-Neurofilament M (BioLegend Cat# 822701), 1:100 fluorescein phalloidin (Thermo Fisher Scientific Cat# F432), 1:500 rabbit anti-MAG (Invitrogen Cat#34-6200), 1:300 goat anti-choline acetyltransferase (Millipore Cat# AB144P), 1:2 mouse anti-Cnx32 (7C6.C7) (a gift from Dr. Steven Scherer[84]), and 1:2,000 mouse anti-tubulin b3 (TuJ1) (Covance Cat# MMS-435P). After washing three times with 1× PBS, the following secondary antibodies were diluted in blocking buffer and applied to sections for 1 h at room temperature: 1:500 Alexa 488 donkey anti-rabbit IgG (Jackson ImmunoResearch Cat# 711-545-152), 1:500 rhodamine (TRITC) donkey anti-rabbit IgG (Jackson ImmunoResearch Cat# 711-025-152), 1:500 Cy3 donkey anti-mouse IgG (Jackson ImmunoResearch Cat# 715-165-150), 1:500 Cy3 donkey anti-chicken IgY (Jackson ImmunoResearch Cat# 703-165-155), 1:500 Alexa 488 donkey anti-chicken IgY (Jackson ImmunoResearch Cat# 703-545-155), and 1:500 Cy3-AffiniPure donkey anti-goat IgG antibody (Jackson ImmunoResearch Cat# 705-165-003). DAPI was added for 5 min at room temperature. Slides were then washed three times with 1× PBS and mounted in Vectashield mounting medium. Staining for Cnx32 (including Kv1.1) was similar to the general protocol, but sciatic nerves were not fixed before teasing. For analysis of Thy1-YFP, sciatic nerves were dissected, fixed in 4% paraformaldehyde for 1 h, and washed with 1× PBS. After a 10-min incubation in 0.1% Triton X-100 in 1× PBS, nerves were washed with 1× PBS, and the perineurium was carefully removed using a 27-gauge needle. Finally, tissue was whole mounted for visualization. Images were acquired with a confocal Leica SP5II or a Zeiss (Oberkochen, Germany) Apotome microscope. Images were analyzed using ImageJ Fiji v1.52p. Quantification of mitochondrial size was performed as described by Della-Flora Nunes *et al*[85], using a single stack of the axonal plane. Corrected total cell fluorescence was calculated as the integrated density – (area of selected cell × mean fluorescence of background readings).

### RNA isolation

Total RNA was isolated from mouse sciatic nerves using TRIzol (Life Technologies, Carlsbad, CA) reagent according to the manufacturer's instructions. Samples were frozen in liquid $N_2$ following dissection and stored at –80°C. A cooled pestle was used to pulverize tissues to powder. The powder was then resuspended in TRIzol reagent. After incubation for 5 min on ice, chloroform was added and the mixture was shaken vigorously and centrifuged. The upper aqueous phase containing RNA was collected, and 1 μL glycogen was added (20 mg/mL). RNA was precipitated with isopropanol, pelleted, and washed twice with 75% ethanol. Pellets were resuspended in 10 μL DEPC-d$H_2O$. RNA was quantified (optical density at 260 nm) using a spectrophotometer (NanoDrop 2000C; Thermo Fisher Scientific) and analyzed for purity according to the 260/280 ratio.

### cDNA preparation and TaqMan qRT-PCR

An Invitrogen kit (Superscript III) was used to convert 1 μg of total RNA to cDNA using the oligo(dT) and hexamers provided. Following the reverse transcription reaction, the provided RNase H was added to each sample. cDNA was collected and stored at −80°C. For RT-qPCR, TaqMan systems were used for different primers. The RT-qPCR reaction was 50°C for 3 min, 95°C for 10 min, 95°C for 15 s, and 60°C for 1 min (40 cycles). The amount of cDNA used in qRT-PCR reactions was determined by standard curves in accordance with Applied Biosystems protocols. Each cDNA sample was tested in triplicates for the presence of each gene of interest and for the standard (*GAPDH*) on the same plates. Target and reference gene PCR amplification was performed with Assays-on-Demand (Applied Biosystems Instruments): *GAPDH* (Mm99999915_g1), *Ddit3*/*Chop* (Mm00492097_m1), *XBP1s* (Mm03464496_m1) and *Hspa5*/*BIP* (Hs99999174_m1). All samples were analyzed in triplicates, and the relative expression of the target RNAs was calculated using the $\Delta\Delta C_T$ of the gene of interest compared to the housekeeper gene. Reactions without target cDNA were used as a negative control for each reaction. Sciatic nerves from 3 to 6 mice per genotype per time point were analyzed.

### Protein extraction and Western blotting

Tissue or cells were lysed in RIPA buffer supplemented with phosphatase and protease inhibitors. For sciatic nerves, the samples were flash frozen in liquid $N_2$ and pulverized with a pestle. The crushed powder was resuspended in lysis buffer. For cells, wells were washed with 1× PBS, and RIPA buffer was added to each well on ice. Cells were scraped off in RIPA buffer and collected. Lysates were left on ice for 20 min and centrifuged at 13,000 × *g* for 20 min at 4°C. Supernatant protein concentrations were determined with a BCA protein assay kit (Thermo Fisher Scientific) according to the manufacturer's instructions. For deglycosylation experiments, samples were treated with endoglycosidase H (New England BioLabs Cat# P0702) and PNGase F (New England BioLabs Cat# P0704) according to the manufacturer's instructions. Samples were prepared with 4× Laemmli buffer and lysis buffer. Five micrograms of protein was loaded per lane and resolved using SDS-polyacrylamide gel electrophoresis (SDS-PAGE) under denaturing conditions with a mini-Protean II gel electrophoresis apparatus; Precision Plus Standard Protein Dual color (Bio-Rad) was included to enable band size identification. Separated proteins were transferred to a polyvinylidene difluoride blotting membrane in a mini gel transfer tank. Blots were then blocked with 5% BSA in TBS-0.05% Tween 20 for 1 h at room temperature. The blots were incubated overnight at 4°C with the following primary antibodies in 3% BSA in TBS-0.05% Tween 20: 1:5,000 chicken anti-P0 (Aves Lab Cat# PZO), 1:1,000 rat anti-HA high affinity (Roche Cat# 11867423001), 1:3,000 rabbit anti b-tubulin (Novus Cat# NB600-936), 1:1,000 rabbit anti-PMP22 (Sigma Cat# SAB4502217), 1:1,000 rabbit anti-CNPase (Cell Signaling Cat# 5664), 1:1,000 rabbit anti-MAG (Invitrogen Cat# 34-6200), and 1:1,000 rabbit anti-HSP90a (Thermo Fisher Scientific Cat# PA3-013). Membranes were washed in TBS-0.05% Tween 20 three times for 5 min and incubated for 1 h at room temperature with the following horseradish peroxidase-conjugated secondary antibodies: 1:20,000 donkey anti-rabbit IgG(H+L) polyclonal (Novus Cat# NB7185) and peroxidase-AffiniPure donkey anti-chicken IgY antibody (Jackson ImmunoResearch Cat# 703- 035-155). Blots were developed using ECL (GE Healthcare, Chicago, IL) and quantified using Image J software. Sciatic nerves from 5 mice per genotype per time point were analyzed.

### Metabolite extraction and measurement

Tissue metabolites extraction and measurement were performed as described by Sasaki *et al*[55]. Tissues were collected and immediately frozen in liquid nitrogen and stored at −80 °C. Frozen tissues were homogenized by sonication (Branson Sonifier 450, output 2.5, 50% duty cycle, 10–20 s) in 50% MeOH in water (160 µl). Homogenates were centrifuged (13,000 *g*, 10 min, 4 °C) and cleared supernatants were transferred to new tubes. One third volume of chloroform was added to the supernatant, mixed, and centrifuged (13,000 *g*, 10 min, 4 °C). The aqueous phase (140 µl) was transferred to a new tube and lyophilized and stored at −20 °C until analysis. Lyophilized samples were reconstituted with 5 mM ammonium formate (70 µl for the sciatic nerve), centrifuged (13,000 *g*, 10 min, 4 °C). The 10 µl clear supernatant was mixed with 10 µl 5mM ammonium formate and loaded on LC-MS.

NAD+, NADH, ATP, ADPR, and NAM were measured using LC-MS/MS using C18 reverse phase column (Atlantis T3, 2.1 × 150 mm, 3 µm; Waters) equipped with HPLC system (Agilent 1290 LC) at a flow rate of 0.15 ml/min with 5 mM ammonium formate for mobile phase A and 100% methanol for mobile phase B. Metabolites were eluted with gradients of 0–10 min, 0–70% B; 10–15 min, 70% B; 16–20 min, 0% B. The metabolites were detected with a triple quadrupole mass spectrometer (Agilent 6470 MassHunter; Agilent) under positive ESI and multiple reaction monitoring (MRM) mode. Lactate, pyruvate, and G6P were measured using Metabolomics dMRM Database and Method (Agilent) according to the instructions. Serial dilutions of standards for each metabolite in 5 mM ammonium formate were used for calibration of NAD+, NADH, ATP, ADPR, and NAM. Metabolites were quantified by MassHunter quantitative analysis tool (Agilent) with standard curves (NAD+, NADH, ATP, ADPR, and NAM) or area under curves (lactate, pyruvate, and G6P) and normalized by the protein amount measured by BCA protein assay kit (Pierce).

### Auditory brainstem response (ABR)

The ABR was recorded in a sound attenuating chamber using a commercial system (SmartEP, Intelligent Hearing Systems, Miami, FL). Mice were anesthetized with ketamine (50 mg/kg, i.p.) and xylazine (6 mg/kg), placed on a temperature-controlled heating pad, and needle electrodes placed on the vertex (non-inverting), behind the ipsilateral pinna (inverting electrode) and the behind the contralateral pinna (ground). Neural responses were amplified, filtered (30 - 3000 Hz) and digitized (1024 presentations, 40 kHz sampling rate) in response to the click stimuli (1 ms rise/fall, cosine gated, 5 ms duration, 21/s) over a 10 ms window. The ABR was elicited with click stimuli to obtain well-defined peaks of wave I to V from which to estimate the latency of wave I. Click intensity was decreased from 90 to 30 dB pSPL in 10 dB steps). A wave I latency versus intensity function was generated for each WT (n=5) and TM/TM (n=5) mouse and the data used to construct a mean wave I latency-intensity function for WT and TM/TM mice.

### Cochleograms

Cytocochleograms were prepared as described previously[86,87]. Mice were killed with an overdose of $CO_2$, decapitated, the temporal bones quickly removed, the round and oval windows opened, and 10% formalin in phosphate buffered saline perfused into the cochlea. The cochleae were immersed in 10% formalin for 24 h, decalcified with 10% EDTA and stained with Harris' hematoxylin solution. The cochlear basilar membrane was dissected out, mounted as a flat surface preparation in glycerin on a glass slide, examined with a light microscope (400X) and the numbers of missing inner hair cells (IHCs) and outer hair cells (OHCs) counted along the entire length of the cochlea. Cochleograms were prepared showing the percentages of missing OHCs and IHCs as a function of percent distance from the apex were generated for each animal. Mean cochleograms were prepared for WT (n=4) and TM/TM (n=5) mice.
