## Supplementary figures for "MPZ-T124M mouse model replicates human axonopathy and suggest alteration in axo-glia communication"

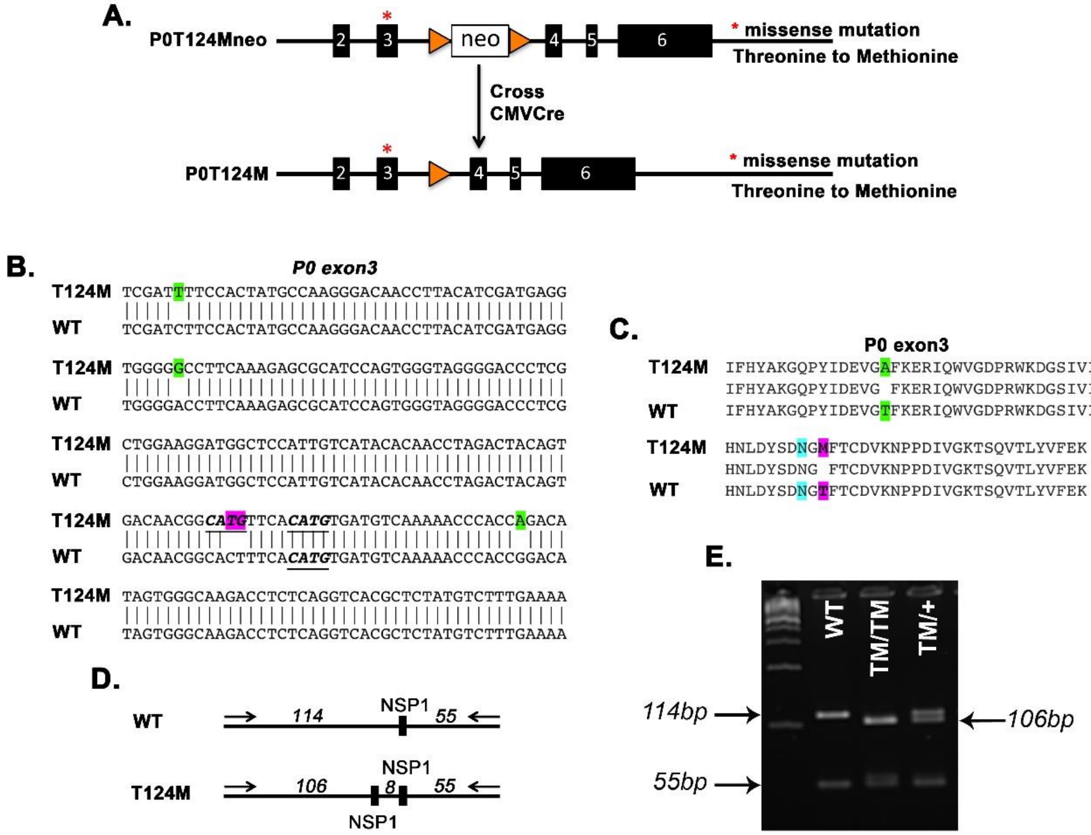


**Figure S1: TM mouse construction and sequence validation**

**Supplementary Figure 1: TM mouse construction and sequence validation.** (**A**) The targeting construct to introduce the T124M mutation into exon 3 of the mouse *Mpz* gene is based on the construct used to engineer the R98C mouse[17]. The neomycin resistance gene was removed by crossing the founder mice with CMV-Cre mice. (**B and C**) P0 mRNA from wild-type (WT) and P0T124M homozygous (T124M) sciatic nerves was cloned and sequenced. Nucleotide (**B**) and amino acid (**C**) sequences (P0 exon 3) for the wild type (WT) and mutant (T124M) were compared. T124M substitution is highlighted in purple. Note how close the T124M mutation is to the *N*-glycosylation acceptor site N122 (highlighted in blue). Strain-specific neutral mutation is indicated in green. More details are provided in the supplemental sequencing file. (**D**) P0T124M mouse genotyping strategy. Introduction of T124M mutation generates a restriction recognition site for NSP1. This new site is located 8 bp from the 5′ end of a preexisting NSP1 restriction site. After amplification, DNA from the wild type (WT) is cut once by NSP1 to generate two fragments of 114 and 55 bp. DNA from the T124M mutant is cut at two different sites by NSP1 to generate three fragments of 106, 8, and 55 bp. (**E**) Representative genotyping PCR of WT, T124M heterozygous, and homozygous mice. WT has a band at 114 bp. T124M heterozygote exhibits the 114 bp band and the 106-bp mutant band. T124M homozygote shows only one band at 106 bp. The three genotypes have a band at 55 bp; 8 bp is too small to be detected.

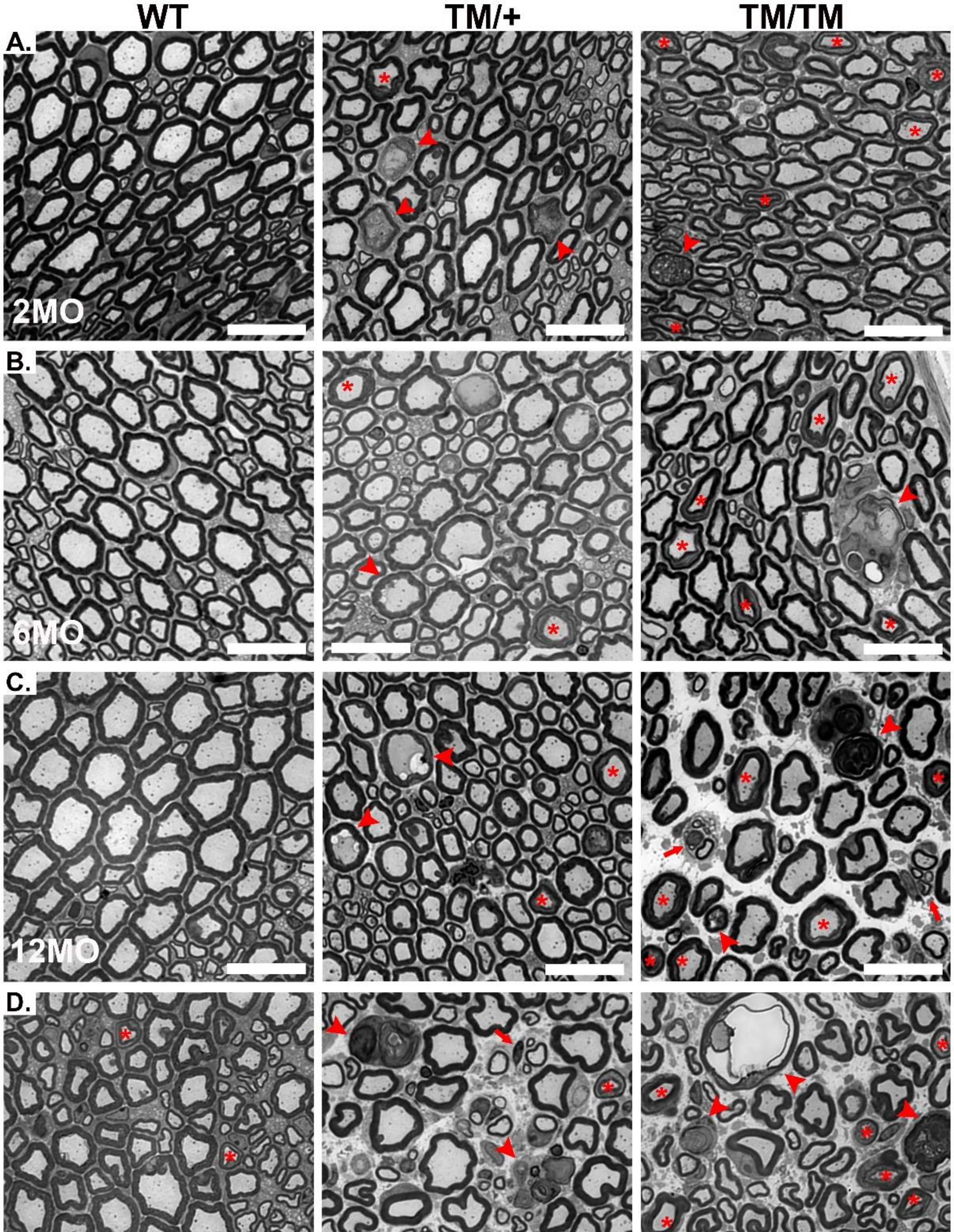


**Figure S2: Semithin transections of sciatic nerves reveal axonal degeneration in TM mice**

**Supplementary Figure 2: Semithin transections of sciatic nerves reveal axonal degeneration in TM mice.** Representative images of transverse semithin sections of sciatic nerves stained with toluidine blue from wild-type (WT) and P0T124M heterozygous (TM/+) and homozygous (TM/TM) mice at 2 (**A**), 6 (**B**), 12 (**C**), and 18 (**D**) months of age. Arrowheads indicate degenerative figures, arrows indicate regenerative axons, asterisks indicate Schmidt-Lanterman incisures (SLI), and “ba” indicates a myelin balloon. Scale bars: 20 μm.

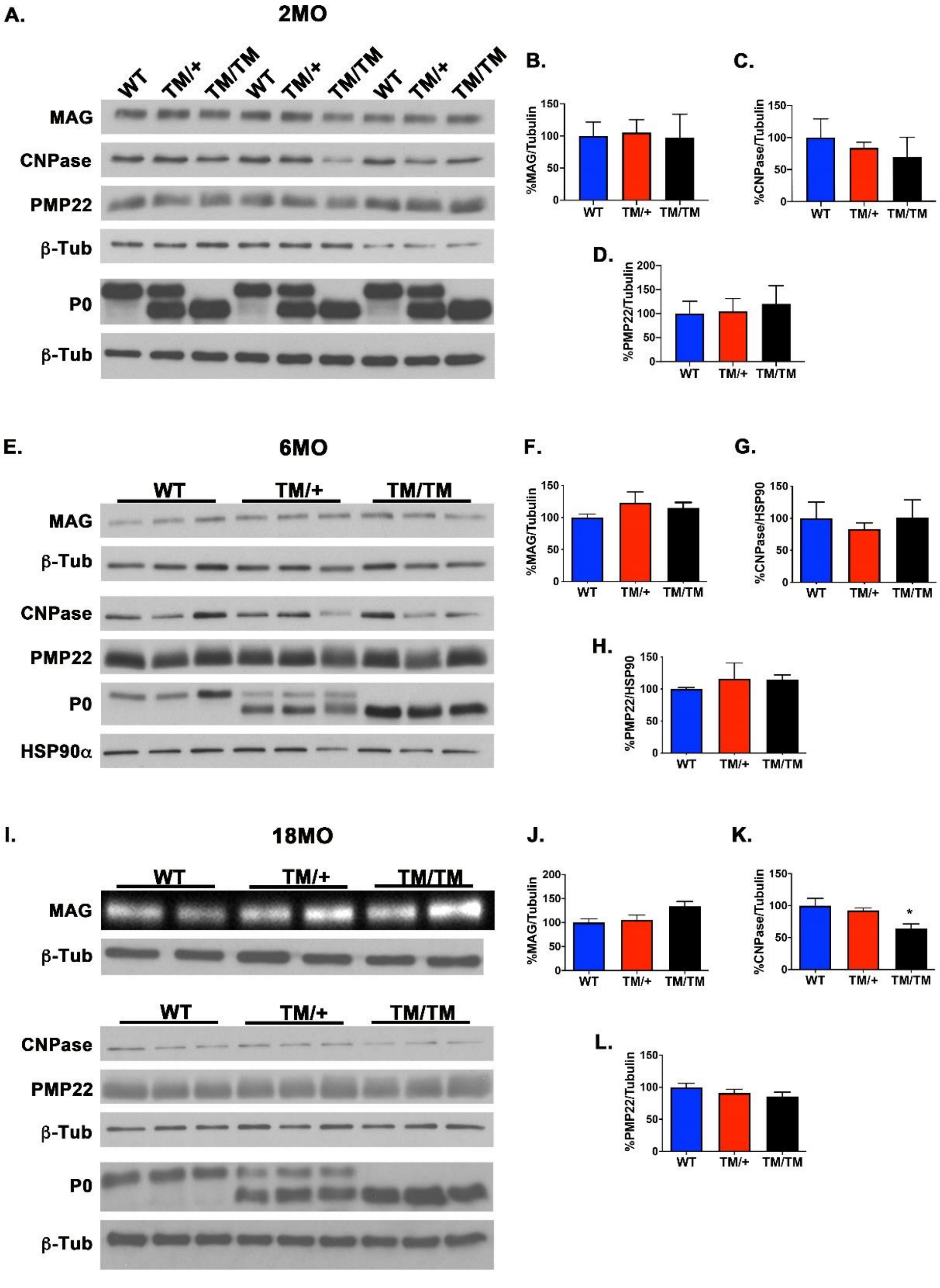


**Figure S3: Myelin proteins expression is not altered in TM mice**

**Supplementary Figure 3: Myelin proteins expression is not altered in TM mice.** Western blot analysis for wild-type (WT) and P0T124M heterozygous (TM/+) and homozygous (TM/TM) mice at 2 (**A**), 6 (**E**), and 18 (**I**) months of age. Blots were probed with CNP, PMP22, and P0 antibodies. β-Tubulin and HSP90α were used as loading controls. Densitometric quantification did not reveal an alteration of MAG (**B, F, and J**) or PMP22 (**D, H, and L**) expression. CNP expression was reduced in TM/TM mice at 18 months of age (**K**) but not at younger ages (**C and G**). T124M mutation alters P0 migration. *n* (animals) = 5 per genotype. $*p < 0.05$ by multiple-comparisons Tukey's *post hoc* tests after one-way ANOVA (**B** to **D**, **F** to **H**, and **J** to **L**). Graphs indicate means ± SEMs.

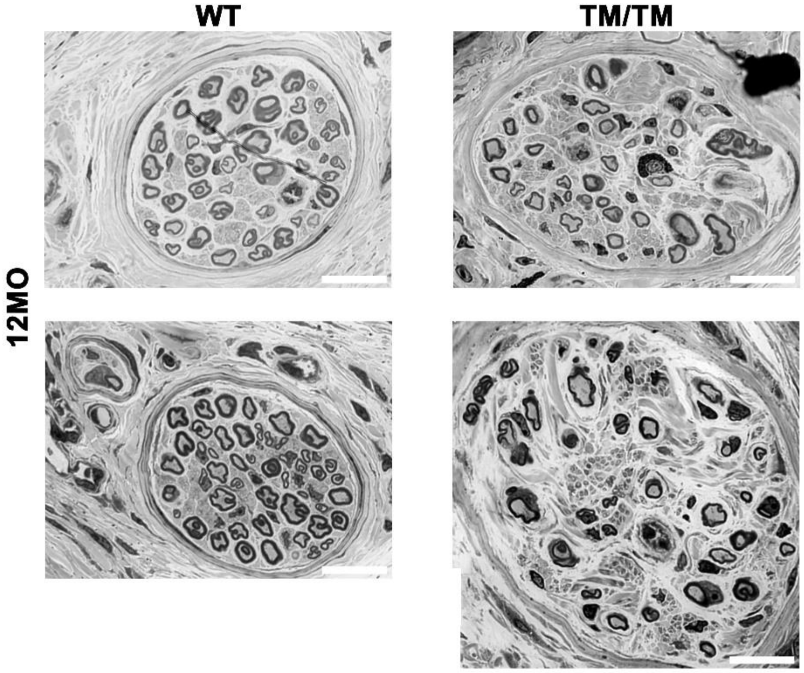


**Figure S4: Semithin transections of digital nerves from toes reveal fiber degeneration in TM mice with an FVB/N background**

**Supplementary Figure 4: Semithin transections of digital nerves from toes reveal fiber degeneration in TM mice with an FVB/N background.** Representative images of transverse semithin sections of digital nerves stained with toluidine blue from two wild-type (WT) and two P0T124M homozygous (TM/TM) mice at 12 months of age in FVB/N background. Scale bars: 20 μm.

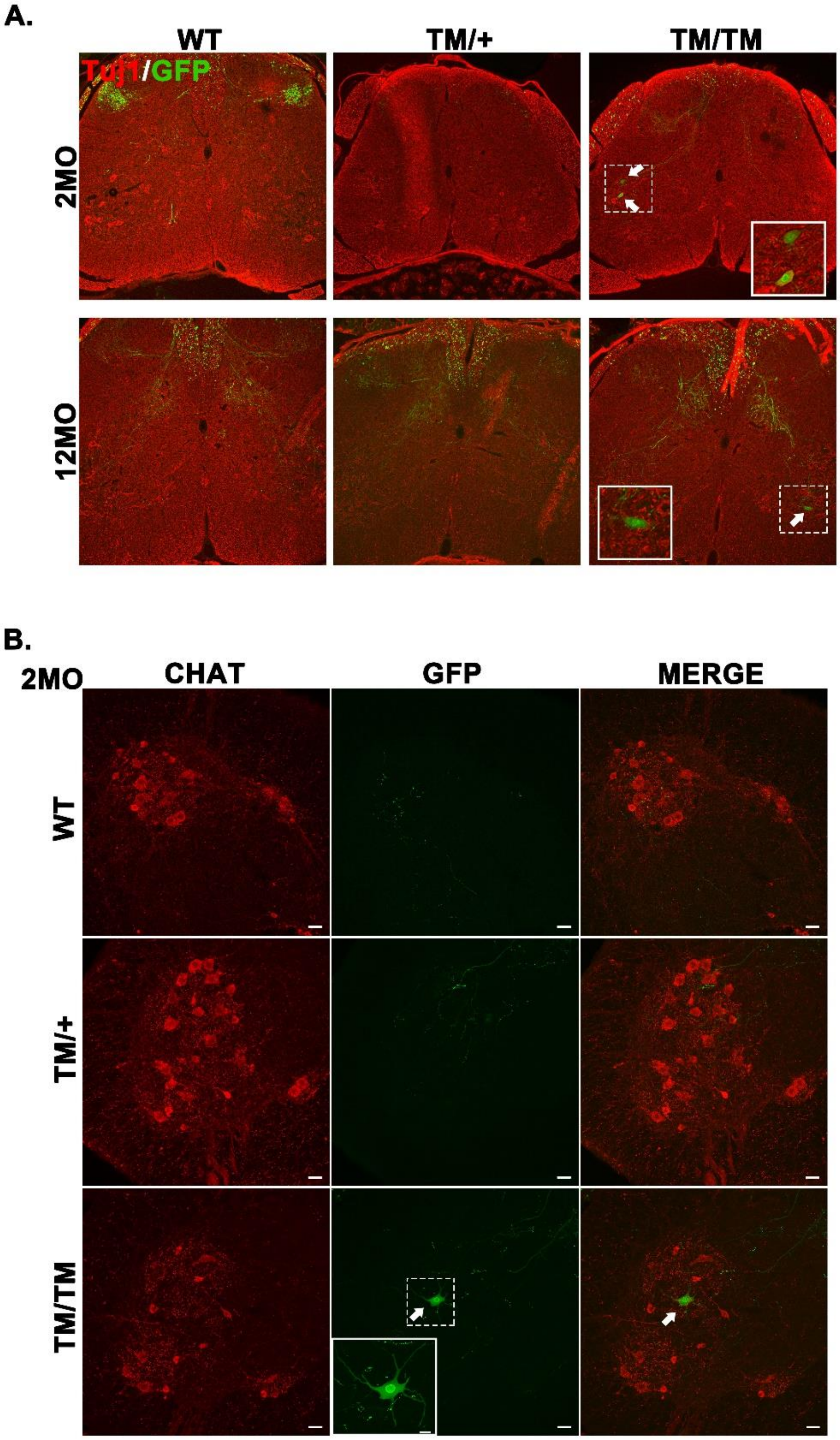


**Figure S5: ATF3 expression in TM spinal cord**

**Supplementary Figure 5: ATF3 expression in TM spinal cord.** Representative confocal microscopy images of spinal cords sections (L3-L5) from 2 and 12-month-old WT, TM/+ and TM/TM–ATF3-GFP mice stained for TuJ1 (red) (**A**), choline acetyltransferase (CHAT; motoneurons) (red) (**B**) and ATF3-GFP (green). Scale bars: 40 μm. High-magnification insets show motoneuron expressing GFP under ATF3 promoter control.

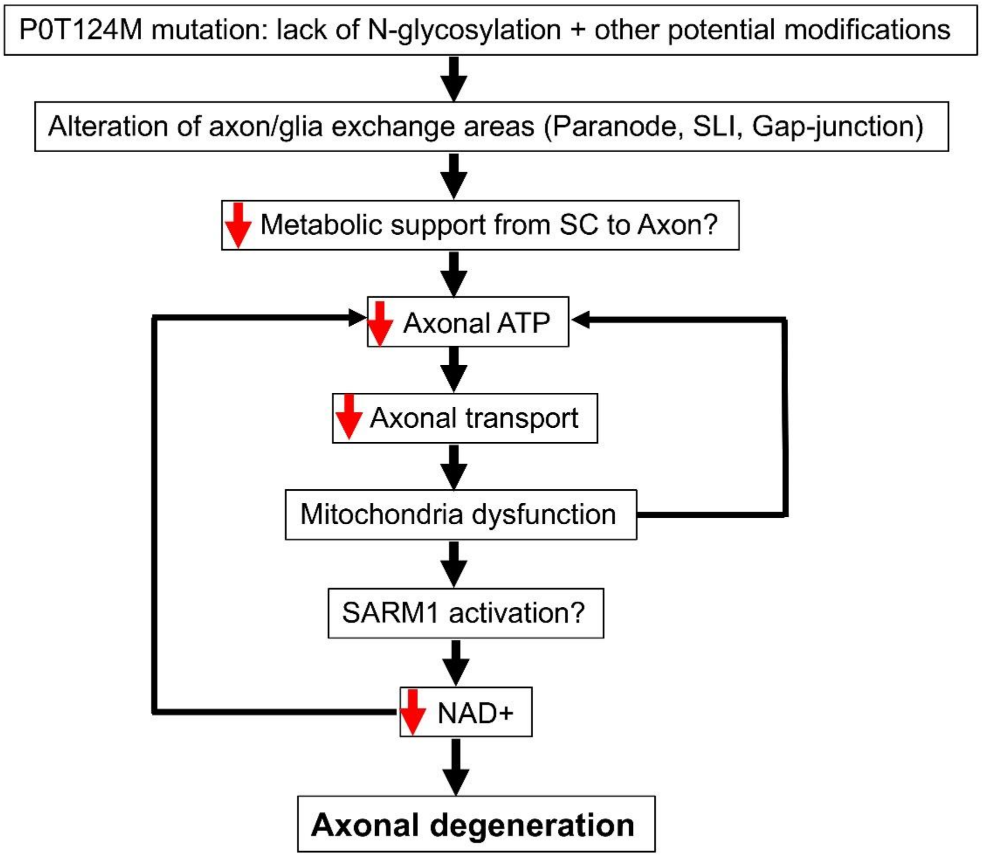


**Figure S6: Hypothetical P0T124M pathomechanism**

**Supplementary Figure 6: Schematic hypothetical P0T124M pathomechanism.** P0T124M mutation impedes *N*-glycosylation and is responsible for additional P0 modifications. Axon–glia exchange areas (paranodes, Schmidt-Lanterman incisures [SLI], and gap junctions) are altered in P0T124M mutants and could lead to deficient transport of metabolites from Schwann cells (SC) to axons. Lack of SC support deprives axons of ATP. Axonal transport is slowed, leading to mitochondrial fragmentation, aggregation, and degeneration. Damaged mitochondria are not able to produce ATP, inducing a vicious cycle. Mitochondrial stress could activate SARM1, which cleaves $NAD^+$. $NAD^+$ depletion leads to axonal degeneration.

**Supplementary Movie 1:** Example of 6-month-old WT mouse performing beam walking test. Related to Figure 1.

**Supplementary Movie 2:** Example of 6-month-old TM/+ mouse performing beam walking test. Related to Figure 1.

**Supplementary Movie 3:** Example of 6-month-old TM/TM mouse performing beam walking test. Related to Figure 1.
