## Supplementary material for "MPZ-T124M mouse model replicates human axonopathy and suggest alteration in axo-glia communication": sequencing

### Sciatic nerve 32717 clone 1 nucleotide sequence

TTTCCTTAGATCATGCTCGAGCGGCCGCCAGTGTGATGGATATCTGCAGAATTCGGCTTATGGCTCCCGGGGCTCC
CTCCTCCAGCCCCAGCCCTATCCTGGCTGCCCTGCTCTTCTCTTCTTTGGTGCTGTCTCCAGCCCTGGCCATTGTGGTT
TACACGGACAGGGAAATCTATGGTGCTGTGGGCTCCCAGGTGACCCTGCACTGCTCCTTCTGGTCCAGTGAATGGG
TCTCAGATGACATCTCTTTTACCTGGCGCTACCAGCCTGAAGGGGGCCGAGATGCCATTTCGATTTTCCACTATGCC
AAGGGACAACCTTACATCGATGAGGTGGGGGCCTTCAAAGAGCGCATCCAGTGGGTAGGGGACCCTCGCTGGAA
GGATGGCTCCATTGTCATACACAACCTAGACTACAGTGACAACGGCATGTTCACATGTGATGTCAAAAACCCACCA
GACATAGTGGGCAAGACCTCTCAGGTCACGCTCTATGTCTTTGAAAAAGTGCCCACTAGGTATGGGGTGGTGTTGG
GAGCAGTGATCGGGGGCATCCTCGGGGTGGTGCTGTTGCTGCTGTTGCTCTTCTACCTGATTCGGTACTGCTGGCT
GCGCAGGCAGGCTGCCCTGCAGAGAAGGCTCAGTGCCATGGAGAAGGGGAGATTTCACAAATCTTCGAAGGACT
CCTCGAAGCGAGGGCGGCAGACGCCAGTGCTGTATGCCATGCTGGACCACAGCCGAAGCACCAAAGCTGCCAGTG
AGAAGAAATCAAAAGGGCTGGGGGAGTCTCGCAAGGATAAGAAAATAGAAGCCGAATTCCAGCACACTGGCGGC
CGTTACTAGTGGATCCGAGCTCGGTACCAAGCTTGGGCGTAATCATGGTCATAGCTGTTTCCTGTGTGAAATTGTTA
TCCGCTCACAATTCCACACAACATACGAGCCGGAAGCATAAAGTGTAAAGCCTGGGGTGCCTAATGAGTGAGCTA
ACTCACATTAATTTGCGTTGCGCTCACTGCCGCTTTCCAGTCGGAAACCTGTCGTGCCAGCTGCATATGATCGGCCA
CGCGCGGGGAGAGGCGTTGCGTATGGCGCCTCTTCGCTCTCGCTTCACTGAACTCGCTGGCGCTCGGTCGATTCCG
GCCTGCGCGGCCGAGACG

Start codon Stop codon

### 1. Standard nucleotide blast NCBI

Mus musculus myelin protein zero (Mpz), mRNA
Sequence ID: ref|NM_008623.4|Length: 1993Number of Matches: 1
Related Information
Gene-associated gene details
GEO Profiles-microarray expression data
Map Viewer-aligned genomic context
Range 1: 167 to 910GenBankGraphics Next Match Previous Match

Alignment statistics for match #1

| Score | Expect | Identities | Gaps | Strand |
|---|---|---|---|---|
| 1310 bits(1452) | 0.0 | 737/744(99%) | 0/744(0%) | Plus/Plus |

```
Query  1    ATGGCTCCCGGGGCTCCCTCCTCCAGCCCCAGCCCTATCCTGGCTGCCCTGCTCTTCTCT  60
            ||||||||||||||||||||||||||||||||||||||||||||||||||||||||||||
Sbjct  167  ATGGCTCCCGGGGCTCCCTCCTCCAGCCCCAGCCCTATCCTGGCTGCCCTGCTCTTCTCT  226

Query  61   TCTTTGGTGCTGTCTCCAGCCCTGGCCATTGTGGTTTACACGGACAGGGAAATCTATGGT  120
            ||||||||||| ||||||||||||||||||||||||||||||||||||||||||||||||
Sbjct  227  TCTTTGGTGCTCTCTCCAGCCCTGGCCATTGTGGTTTACACGGACAGGGAAATCTATGGT  286

Query  121  GCTGTGGGCTCCCAGGTGACCCTGCACTGCTCCTTCTGGTCCAGTGAATGGGTCTCAGAT  180
            || |||||||||||||||||||||||||||||||||||||||||||||||||||||||||
Sbjct  287  GCCGTGGGCTCCCAGGTGACCCTGCACTGCTCCTTCTGGTCCAGTGAATGGGTCTCAGAT  346

Query  181  GACATCTCTTTTACCTGGCGCTACCAGCCTGAAGGGGGCCGAGATGCCATTTCGATTTTC  240
            |||||||||||||||||||||||||||||||||||||||||||||||||||||||| |||
Sbjct  347  GACATCTCTTTTACCTGGCGCTACCAGCCTGAAGGGGGCCGAGATGCCATTTCGATCTTC  406
```

```
Query  241   CACTATGCCAAGGGACAACCTTACATCGATGAGGTGGGGGCCTTCAAAGAGCGCATCCAG  300
             |||||||||||||||||||||||||||||||||||||||| ||||||||||||||||||||
Sbjct  407   CACTATGCCAAGGGACAACCTTACATCGATGAGGTGGGGACCTTCAAAGAGCGCATCCAG  466

Query  301   TGGGTAGGGGACCCTCGCTGGAAGGATGGCTCCATTGTCATACACAACCTAGACTACAGT  360
             ||||||||||||||||||||||||||||||||||||||||||||||||||||||||||||
Sbjct  467   TGGGTAGGGGACCCTCGCTGGAAGGATGGCTCCATTGTCATACACAACCTAGACTACAGT  526

Query  361   GACAACGGCATGTTCACATGTGATGTCAAAAACCCACCAGACATAGTGGGCAAGACCTCT  420
             ||||||||||  |||||||||||||||||||||||||| |||||||||||||||||||||
Sbjct  527   GACAACGGCACTTTCACATGTGATGTCAAAAACCCACCGGACATAGTGGGCAAGACCTCT  586

Query  421   CAGGTCACGCTCTATGTCTTTGAAAAAGTGCCCACTAGGTATGGGGTGGTGTTGGGAGCA  480
             ||||||||||||||||||||||||||||||||||||||||||||||||||||||||||||
Sbjct  587   CAGGTCACGCTCTATGTCTTTGAAAAAGTGCCCACTAGGTATGGGGTGGTGTTGGGAGCA  646

Query  481   GTGATCGGGGGCATCCTCGGGGTGGTGCTGTTGCTGCTGTTGCTCTTCTACCTGATTCGG  540
             ||||||||||||||||||||||||||||||||||||||||||||||||||||||||||||
Sbjct  647   GTGATCGGGGGCATCCTCGGGGTGGTGCTGTTGCTGCTGTTGCTCTTCTACCTGATTCGG  706

Query  541   TACTGCTGGCTGCGCAGGCAGGCTGCCCTGCAGAGAAGGCTCAGTGCCATGGAGAAGGGG  600
             ||||||||||||||||||||||||||||||||||||||||||||||||||||||||||||
Sbjct  707   TACTGCTGGCTGCGCAGGCAGGCTGCCCTGCAGAGAAGGCTCAGTGCCATGGAGAAGGGG  766

Query  601   AGATTTCACAAATCTTCGAAGGACTCCTCGAAGCGAGGGCGGCAGACGCCAGTGCTGTAT  660
             ||||||||||||||||||||||||||||||||||||||||||||||||||||||||||||
Sbjct  767   AGATTTCACAAATCTTCGAAGGACTCCTCGAAGCGAGGGCGGCAGACGCCAGTGCTGTAT  826

Query  661   GCCATGCTGGACCACAGCCGAAGCACCAAAGCTGCCAGTGAGAAGAAATCAAAAGGGCTG  720
             ||||||||||||||||||||||||||||||||||||||||||||||||||||||||||||
Sbjct  827   GCCATGCTGGACCACAGCCGAAGCACCAAAGCTGCCAGTGAGAAGAAATCAAAAGGGCTG  886

Query  721   GGGGAGTCTCGCAAGGATAAGAAA  744
             ||||||||||||||||||||||||
Sbjct  887   GGGGAGTCTCGCAAGGATAAGAAA  910
```

Neutral mutation du to strain specificity   T124M mutation **Exon3**

**<u>Amino Acid sequence translation from 32717-clone1 nucleotide sequence</u>**

MAPGAPSSSPSPILAALLFSSLVLSPALAIVVYTDREIYGAVGSQVTLHCSFWSSEWVSDDISFTWRYQPEGGRDAISIFHY
AKGQPYIDEVGAFKERIQWVGDPRWKDGSIVIHNLDYSDNGMFTCDVKNPPDIVGKTSQVTLYVFEKVPTRYGVVLGA
VIGGILGVVLLLLLLFYLIRYCWLRRQAALQRRLSAMEKGRFHKSSKDSSKRGRQTPVLYAMLDHSRSTKAASEKKSKGLG
ESRKDKK

**<u>Amino Acid sequence from P0 *mus musculus* C57B6 NM_008623</u>**

MAPGAPSSSPSPILAALLFSSLVLSPALAIVVYTDREIYGAVGSQVTLHCSFWSSEWVSDDISFTWRYQPEGGRDAISIFHY
AKGQPYIDEVGTFKERIQWVGDPRWKDGSIVIHNLDYSDNGTFTCDVKNPPDIVGKTSQVTLYVFEKVPTRYGVVLGAV
IGGILGVVLLLLLLFYLIRYCWLRRQAALQRRLSAMEKGRFHKSSKDSSKRGRQTPVLYAMLDHSRSTKAASEKKSKGLGE
SRKDKK

### 2. Emboss water protein alignment

Aligned_sequences: 2 # 1: EMBOSS_001 # 2: EMBOSS_001 # Matrix: EBLOSUM62 # Gap_penalty: 10.0 # Extend_penalty: 0.5 # # Length: 248 # Identity:246/248 (99.2%) # Similarity: 246/248 (99.2%) # Gaps: 0/248 ( 0.0%) # Score: 1283.0

```
MAPGAPSSSPSPILAALLFSSLVLSPALAIVVYTDREIYGAVGSQVTLHC
||||||||||||||||||||||||||||||||||||||||||||||||||
MAPGAPSSSPSPILAALLFSSLVLSPALAIVVYTDREIYGAVGSQVTLHC
SFWSSEWVSDDISFTWRYQPEGGRDAISIFHYAKGQPYIDEVGAFKERIQ
|||||||||||||||||||||||||||||||||||||||||||.||||||
SFWSSEWVSDDISFTWRYQPEGGRDAISIFHYAKGQPYIDEVGTFKERIQ
WVGDPRWKDGSIVIHNLDYSDNGMFTCDVKNPPDIVGKTSQVTLYVFEKV
|||||||||||||||||||||||.||||||||||||||||||||||||||
WVGDPRWKDGSIVIHNLDYSDNGTFTCDVKNPPDIVGKTSQVTLYVFEKV
PTRYGVVLGAVIGGILGVVLLLLLLFYLIRYCWLRRQAALQRRLSAMEKG
||||||||||||||||||||||||||||||||||||||||||||||||||
PTRYGVVLGAVIGGILGVVLLLLLLFYLIRYCWLRRQAALQRRLSAMEKG
RFHKSSKDSSKRGRQTPVLYAMLDHSRSTKAASEKKSKGLGESRKDKK
||||||||||||||||||||||||||||||||||||||||||||||||
RFHKSSKDSSKRGRQTPVLYAMLDHSRSTKAASEKKSKGLGESRKDKK
```

Neutral mutation du to strain specificity T124M mutation

### 3. Standard protein blast NCBI

**myelin protein P0 isoform L-MPZ precursor [Mus musculus]**

**Sequence ID: NP_001302429.1Length: 312Number of Matches: 1**

Identities:246/248(99%), Positives: 246/248(99%), Gaps: 0/248(0%)

```
MAPGAPSSSPSPILAALLFSSLVLSPALAIVVYTDREIYGAVGSQVTLHCSFWSSEWVSD
MAPGAPSSSPSPILAALLFSSLVLSPALAIVVYTDREIYGAVGSQVTLHCSFWSSEWVSD
MAPGAPSSSPSPILAALLFSSLVLSPALAIVVYTDREIYGAVGSQVTLHCSFWSSEWVSD

DISFTWRYQPEGGRDAISIFHYAKGQPYIDEVGAFKERIQWVGDPRWKDGSIVIHNLDYS
DISFTWRYQPEGGRDAISIFHYAKGQPYIDEVG FKERIQWVGDPRWKDGSIVIHNLDYS
DISFTWRYQPEGGRDAISIFHYAKGQPYIDEVGTFKERIQWVGDPRWKDGSIVIHNLDYS

DNGMFTCDVKNPPDIVGKTSQVTLYVFEKVPTRYGVVLGAVIGGILGVVLLLLLLFYLIR
DNG FTCDVKNPPDIVGKTSQVTLYVFEKVPTRYGVVLGAVIGGILGVVLLLLLLFYLIR
DNGTFTCDVKNPPDIVGKTSQVTLYVFEKVPTRYGVVLGAVIGGILGVVLLLLLLFYLIR

YCWLRRQAALQRRLSAMEKGRFHKSSKDSSKRGRQTPVLYAMLDHSRSTKAASEKKSKGL
YCWLRRQAALQRRLSAMEKGRFHKSSKDSSKRGRQTPVLYAMLDHSRSTKAASEKKSKGL
YCWLRRQAALQRRLSAMEKGRFHKSSKDSSKRGRQTPVLYAMLDHSRSTKAASEKKSKGL

GESRKDKK
GESRKDKK
GESRKDKK
```

Neutral mutation du to strain specificity T124M mutation
